# Human brain organoid networks

**DOI:** 10.1101/2021.01.28.428643

**Authors:** Tal Sharf, Tjitse van der Molen, Stella M.K. Glasauer, Elmer Guzman, Alessio P. Buccino, Gabriel Luna, Zhouwei Cheng, Morgane Audouard, Kamalini G. Ranasinghe, Kiwamu Kudo, Srikantan S. Nagarajan, Kenneth R. Tovar, Linda R. Petzold, Andreas Hierlemann, Paul K. Hansma, Kenneth S. Kosik

**Affiliations:** Neuroscience Research Institute, University of California Santa Barbara, Santa Barbara, CA 93106, USA.; Department of Molecular, Cellular and Developmental Biology, University of California Santa Barbara, Santa Barbara, CA 93106, USA; Department of Biosystems Science and Engineering, ETH Zürich, Mattenstrasse 26, Basel, 4058 Switzerland; Department of Computer Science, University of California Santa Barbara, Santa Barbara, CA 93106, USA; Memory and Aging Center, Department of Neurology, University of California San Francisco, San Francisco, CA 94158, USA; Department Radiology and Biomedical Imaging, University of California San Francisco, San Francisco, CA 94143, USA; Department of Physics, University of California Santa Barbara, Santa Barbara, CA 93106

**Keywords:** human brain organoids, functional connectivity, extracellular recording, CMOS microelectrode arrays, single-unit activity, network oscillations, local field potentials, theta oscillations, phase-locking

## Abstract

Human brain organoids replicate much of the cellular diversity and developmental anatomy of the human brain. However, the physiological behavior of neuronal circuits within organoids remains relatively under-explored. With high-density CMOS microelectrode arrays (26,400 electrodes) and shank electrodes (960 electrodes), we probed broadband and three-dimensional extracellular field recordings generated by spontaneous activity of human brain organoids. These recordings simultaneously captured local field potentials (LFPs) and single-unit activity extracted through spike sorting. From spiking activity, we estimated a directed functional connectivity graph of synchronous neural network activity, which showed a large number of weak functional connections enmeshed within a network skeleton of significantly fewer strong connections. Treatment of the organoid with a benzodiazepine induced a reproducible signature response that shortened the inter-burst intervals, increased the uniformity of the firing pattern within each burst and decreased the population of weakly connected edges. Simultaneously examining the spontaneous LFPs and their phase alignment to spiking showed that spike bursts were coherent with theta oscillations in the LFPs. Our results demonstrate that human brain organoids have self-organized neuronal assemblies of sufficient size, cellular orientation, and functional connectivity to co-activate and generate field potentials from their collective transmembrane currents that phase-lock to spiking activity. These results point to the potential of brain organoids for the study of neuropsychiatric diseases, drug mechanisms, and the effects of external stimuli upon neuronal networks.

## Introduction

Under a variety of conditions^1^, 3D assemblies of pluripotent stem cells can differentiate into a wide diversity of brain cell types and assume certain anatomical features that resemble the developing brain as well as some mature features such as dendritic spines, inhibitory and excitatory synapses^2–4^ with presynaptic vesicles^5^. Neurons within these 3D assemblies generate action potentials upon depolarization^6^, display excitatory and inhibitory post synaptic currents^7^ and exhibit spontaneous network activity as measured by calcium imaging^6,8–10^ and by extracellular field potential recordings from a small number of electrodes^5,11–14^. Progress in the development of flexible electronics have also enabled three-dimensional readout of electrical activity across the surface of an organoid^15^, and recent work has extended the activity repertoire of organoids to include rhythmic activity over a range of oscillatory frequencies^10,16^. However, technological limitations have restricted broadband detection of electrical activity to small numbers of neurons. Consequently, technical shortcomings have hindered attempts to determine functional connectivity among large numbers of neurons, how they collectively synchronize with rhythmic activity and how pharmacologic perturbations affect these physiologic parameters.

Recent advances in CMOS-based microelectrode array (MEA) technologies^17^ have enabled large-scale *ex vivo* mapping of cortical synaptic projections^18^. We utilize this state-of-the-art technology to generate detailed electrical activity maps from human brain organoids using arrays with 26,400 recording electrodes that tile across ≈ 8.01 mm^2^ surface area equivalent to the cross-section of our brain organoids^19^. The densely tiled recording electrodes (electrode diameter of 7.5 μm at a center-to-center distance of 17.5 μm) are spaced at approximately the size of a neuronal soma, and therefore enable robust and accurate assignment of single-unit activity by utilizing electrode redundancy and the characteristic waveform shapes determined by the neuron’s location relative to the recording electrode^20,21^. These technological advances offer a significant improvement over the more limited ascertainment of multi-unit activity obtained from considerably sparser arrays used in previous studies. In addition, we utilized a 960 electrode Neuropixels CMOS shank probe^22^ with an electrode pitch (≈ 20 μm) approximately the size of a neuron soma, to gain access to the z-plane of the organoid. Utilizing the sub-millisecond temporal resolution of extracellular action potential recordings, we quantified the non-linear probability distribution of firing rates across the organoid. A benzodiazepine shifted the firing rate probability distribution curve toward lower firing rate modes. These features were accompanied by more stereotyped population-level burst dynamics. We next constructed a spatial connectivity map derived from pairwise correlations between spiking units measured by the thousand microelectrodes selected from the total set of 26,400 electrodes with the highest activity. A subnetwork of single-unit spikes was identified with a marked increase in pairwise spike correlation strength when modulated pharmacologically with diazepam. We detected theta frequency oscillations supported by their coherence across the surface of the organoid and stereotypical sets of neuronal ensembles phase-locked to the theta oscillation. These results enhance the utility of brain organoids for novel approaches to the study of the neuropharmacological underpinning of network behavior, as well as conditions such as schizophrenia, autism and neurodegeneration^23,24^.

## Results

### High resolution spike map across a human brain organoid

500 μm thick organoid slices were positioned over the recording electrode surface of high-density CMOS MEAs (MaxOne, Maxwell Biosystems, Zurich, Switzerland) and seated with a sterile custom harp slice grid for recording up to six months after placement on the arrays (Supplementary Fig. 1, Methods). Slicing of organoids can enhance cell growth, prevent interior cell death^12^ and promote the development of extensive axon outgrowth^11^. In this study we present electrical activity from six organoids positioned on 2D CMOS arrays and performed pharmacological manipulations on a subset (*n* = 4) of that cohort. Acute electrophysiology measurements were performed on an additional three intact whole organoids using CMOS shank electrodes. Once positioned on the MEA, spiking activity occurred within approximately two weeks across organoids, followed by increasing firing rates in the form of synchronized bursts at about six months (Supplementary Table 1) and were maximally active at about seven months (Supplementary Fig. 2a). Strong correlations between spike pairs emerged that were interpreted as putative connectivity (Supplementary Fig. 2b, see section on short-term interactions). A high-density field recording was obtained from tissue areas of approximately 2.1 mm by 3.85 mm. A spike-activity map was derived by performing an automated scan of contiguous tiled blocks selected from 26,400 recording electrode sites using switch-matrix CMOS technology^19,25^ that enabled configurable routing of up to 1,024 simultaneous recording electrode sites across the surface of the array to on-chip readout electronics (Fig. 1a). Selecting the top 1,020 electrodes based on spiking activity for simultaneous recording, we constructed a spatial map of extracellular action potential waveforms generated by single-unit activity (Fig. 1b,c) as determined by the spike-sorting algorithm Kilosort2 ^20^, which has optimal accuracy and precision based on ground truth data for arrays with similar electrode densities ^21^. Utilizing the single-unit spike times we generated detailed spatiotemporal maps of spiking activity often in the form of synchronized bursts (Fig. 1d) composed of sets of units from across the tissue section (Fig. 1a). To validate that spikes arose from fast synaptic transmission, we blocked AMPA and NMDA receptors with bath application of NBQX (10 μM) and R-CPP (20 μM) and GABA_A_ receptors with gabazine (10 μM) resulting in a 72% ± 29% (mean ± STD) reduction in spiking (Supplementary Fig. 3; *n* = 4 organoids). The remaining action potentials likely reflect interneurons firing in the absence of synaptic input ^26^. A saturating concentration of the sodium-channel blocker tetrodotoxin (1 μM) was subsequently applied to the same organoid slices, resulting in a 98% ± 1% (mean ± STD) reduction in spiking activity compared to control conditions (Supplementary Fig. 3), indicating that falsely detected spikes are a small fraction of the detected signals.

**Figure 1.**
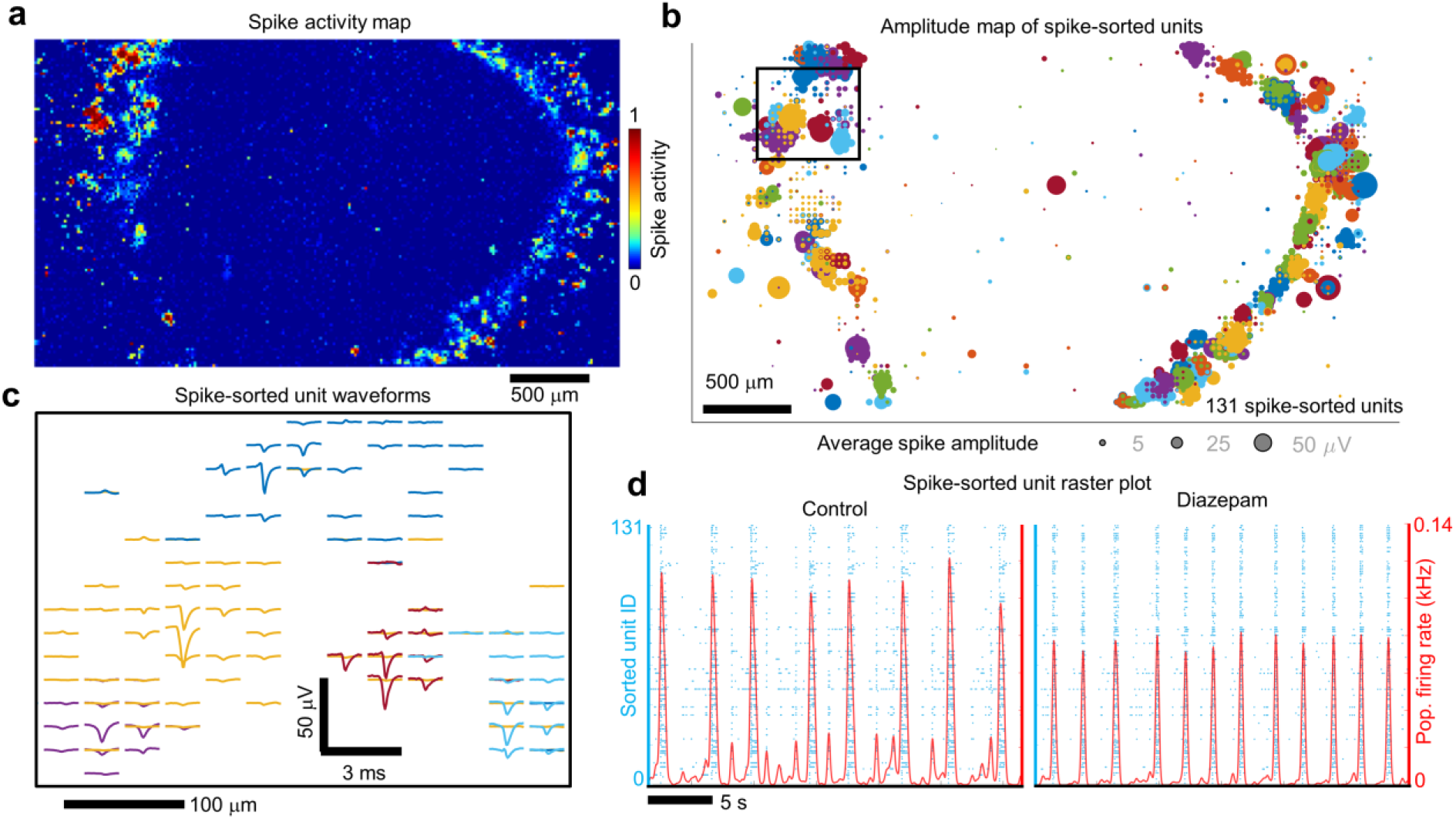
High-resolution maps of extracellular action potentials across a human cerebral organoid. **a** Spatial map of extracellular action potential activity recorded from a 500 μm thick human cerebral organoid slice positioned on a high-density CMOS microelectrode array with 26,400 recording electrodes to survey electrical activity across the entire organoid. The color scale indicates the normalized number of detected spikes (above 5x-rms noise) registered at each electrode site measured over a 30 s interval. Scale bar 500 μm. **b** Spatial map of the mean extracellular action potential spike amplitude (bubble size) from single-unit activity measured simultaneously across the CMOS array from the top 1,020 electrode sites based on activity. A total of 131 spike-sorted units were determined by Kilosort2. Single-unit electrode clusters are plotted using the same color (the same colors are repeated for different units). **C** Extracellular spike waveform traces from four individual units (from the region highlighted by the black square in **b** are plotted with respect to the electrode positions on the array. The sorted unit colors are the same as **b**. Waveform scale bars are 50 μV and 3 ms; spatial scale bar is 100 μm. **d** Left, raster plot of endogenous spiking activity (blue dots) measured from the 131 spike-sorted units. Red line is the average number of spikes detected across the organoid (averaged over a 100 ms Gaussian kernel). Right, same organoid after 50 μM diazepam treatment.

To validate the organoid anatomy as potentially capable of supporting the physiology observed in Figure 1, we characterized the cellular composition with regard to the presence of cell types crucial for the formation, function and maintenance of networked neuronal circuitry ^27,28^. We chose a time points that coincided with the emergence of strong pairwise spike correlations in the organoids. Whole-sections from 8-month old organoids demonstrated the presence of stellate astrocytes expressing gap junction proteins (Fig. 2a). Axons coursed through dendritic fields that appeared directionally aligned in some regions and can project over long distances (Fig. 2b). Multiple interneuron types were present including parvalbumin-labeled cells (Fig. 2c) ^7,29^, as well as synaptobrevin decorated MAP2-positive dendrites (Fig. 2d). Because organoids often have a relatively nutrient deficient center (Fig. 2a), likely due to diffusion limits of the media that reduces cell viability, we suspect that a neuronal poor core accounts for the peripheral ring of higher spiking activity (Fig. 1a and Supplementary Fig. 4 for spiking activity from additional organoids). To obtain additional detail about the cellular composition of the organoids we performed single-cell sequencing on three 7-month-old brain organoids. Unsupervised clustering and dimensionality reduction resulted in the separation of nine major cell populations consisting of cortical excitatory neurons, GABAergic interneurons, astrocytes, an additional VIM+ glial cluster, oligodendrocyte progenitors, and a small cluster of choroid plexus cells (Fig. 2e and Supplementary Fig. 5). GABAergic cells expressed both GAD1 and GAD2 (Supplementary Fig. 6, Supplementary Table 2) as well as several DLX genes indicating their origin in the ventral forebrain (Supplementary Fig. 6). Within both neuronal clusters (glutamatergic and GABAergic), several subunits of GABA receptors associated with network oscillations ^30^ and AMPA and NMDA receptor genes were expressed (Supplementary Fig. 6). The forebrain marker FOXG1 was expressed in all these major cell populations. In contrast, markers for other brain regions (retina, midbrain and hindbrain) were only minimally expressed (Supplementary Fig. 6), indicating a predominantly forebrain identity of our organoids ^31^.

**Figure 2.**
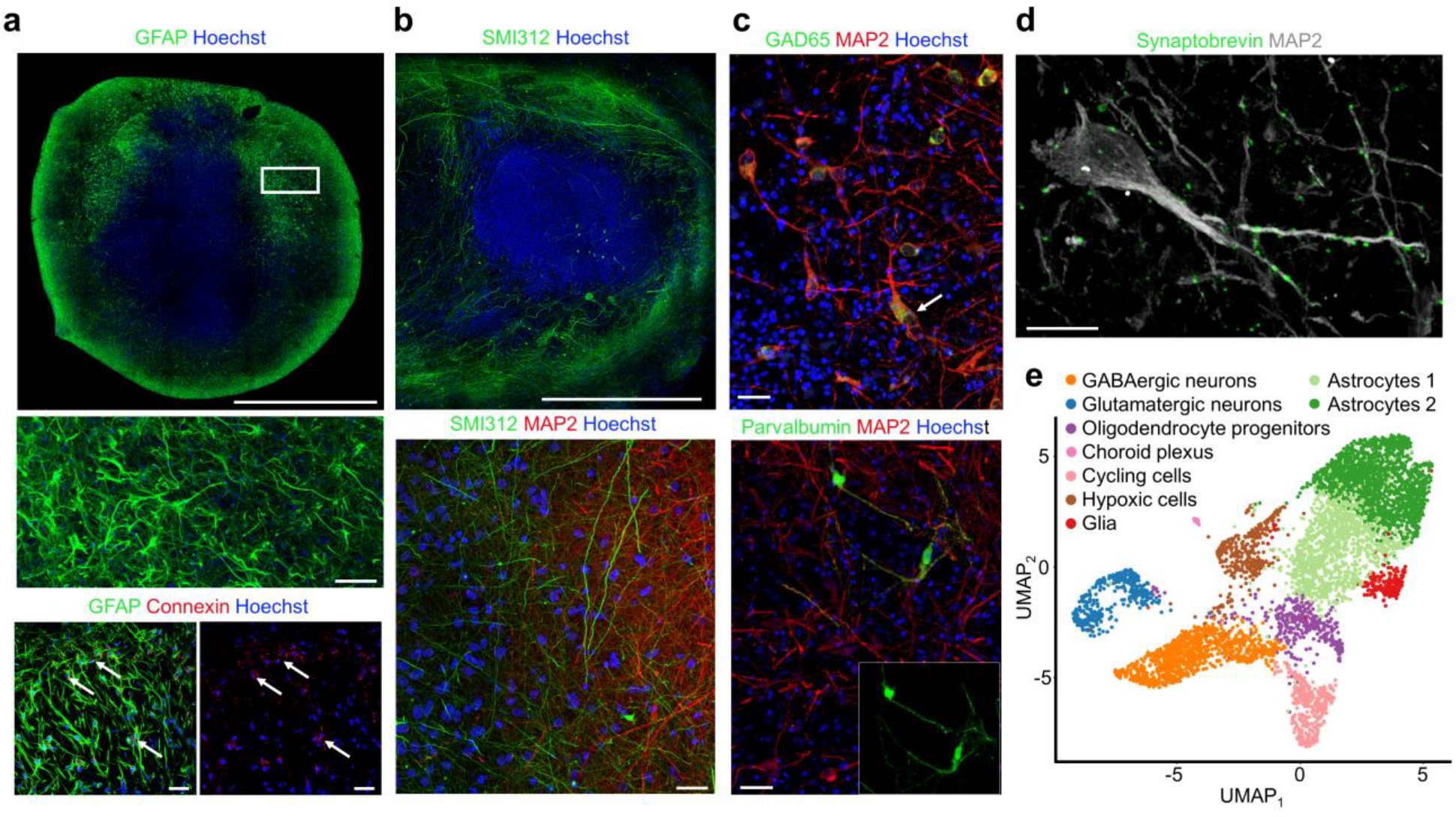
Human cerebral organoids form a scaffolding capable of supporting neuronal microcircuitry. **a** Top, high-resolution, whole-section of an 8-month organoid immunostained with anti-GFAP (green), and counterstained with Hoechst (cell nuclei, blue), scale bar 1 mm. Middle, a high magnification of (top box) showing stellate appearance characteristic of astrocytes. Bottom left, Anti-GFAP-positive astrocytes (green) in an 8-month organoid co-labeled with anti-connexin 43 demonstrating gap junctions (red). Bottom right, connexin 43 gap junction (red; arrows). Middle and bottom scale bars are 40 μm. **b** Top, long neuronal processes labeled with anti-SMI312 (green) in an 8-month organoid, scale bar 500 μm. Bottom, anti-SMI312 (green) axons neighboring MAP2-positive (red) neurons near the margin of 8-month organoid, scale bar 20 μm. **c** Top, anti-GAD65 positive neurons (green) co-labeled with anti-MAP2. Bottom, Anti-Parvalbumin-positive neurons (green) in an 8-month organoid co-labeled with anti-MAP2 (red). Scale bars 40 μm **d** The pre-synaptic marker synaptobrevin label MAP2-positive processes as puncta, scale bar 20 μm. **e** Single-cell RNA sequencing (drop-seq) shows the presence of glutamatergic neurons, GABAergic neurons and astrocyte populations. Single-cell transcriptomes (5,680 cells collected from three 7-month-old organoids) are visualized as a Uniform Manifold Approximation and Projection (UMAP).

Our results expand on and are consistent with previous reports suggesting that self-assembled brain organoids possess the basic architectural motifs necessary to form functional neuronal circuits in 3D space ^2–8,11,12,16^. The resulting circuitry has the capacity to form complex, synaptically coupled electrogenic networks, composed of glutamatergic neurons and GABAergic interneurons which are critically involved in driving network oscillations ^30^. Below, we quantify the precise spatiotemporal signatures of spikes and extracellular field potentials that arise from endogenous and pharmacologically induced activity in brain organoids

### Modulating signaling in brain organoid neuronal circuits with diazepam

Diazepam, a benzodiazepine, is a widespread anxiolytic-hypnotic drug that has a range of therapeutic actions on the central nervous system that range from sedation at low doses to induction of anesthesia at considerably higher doses ^32–34^. In the presence of GABA, diazepam can modulate GABA_A_-receptor channels through two separable mechanisms that result in biphasic potentiation with distinct components in the nM and µM concentration ranges ^34^, with potentiation being very small for the nanomolar range, but significantly larger for the micromolar range (20 µM and higher). We sought to investigate the role diazepam on the electrical properties of neuronal networks in organoids. Diazepam at a concentration of 50 μM decreased the inter-burst intervals, burst duration and the number of spikes participating in the burst (burst amplitude) throughout the organoid (*n* = 4 organoids, Fig. 1d and Supplementary Fig. 8 for diazepam dose responses over the range from 3-50 μM and additional statistics), consistent with diazepam effects in neocortical slice cultures over the same concentration range ^35^.

To quantify the firing rate regimes of single-unit activity, we fit the firing rate probability distributions to a generalized gamma distribution, wherein the shape parameters are equal to *k* and correspond to the Weibull distribution (Fig. 3a, see Methods for additional details). The cumulative Weibull distribution function is of the form *g_λ,k_* (*R*) = 1 – e^−(*R*/*λ*)*k*^, where *R* is the firing rate and *k*, *λ* are the shape and scale parameters, respectively. Recent work *in vivo* has utilized this analytical approach to quantify firing regimes in homologous brain regimes across species ^36^; here we have extended this approach to quantify the pharmacologically induced firing pattern changes in brain organoids. Treatment with diazepam (50 μM) dramatically altered the neuron firing pattern (Fig. 3a), highlighted by the raster plots (bottom panels) for control (blue) and diazepam (red) as well as their respective cumulative distributions (top panels). A two-parameter (λ and *k*) fit to the cumulative firing rate distributions is shown by the dotted lines along with their *R*-squared values for two representative spike-sorted units from organoid L1. Notably, the firing rate regimes are skewed towards higher frequency domains under control conditions relative to diazepam treatment. The cumulative firing rate distributions across the organoid are visualized in Figure 3b for control (blue) and diazepam treatment (red). The upper and lower shaded regions represent the 80^th^ and 20^th^ percentile firing rate bounds across the organoid, highlighting a downshift in the firing rate distribution induced by diazepam. Furthermore, we find that the shape parameter *k* remains similar for control and diazepam treatment (Fig. 3c). Meanwhile, the distribution of scale parameters (λ) for a given organoid exhibited a significant downshift when treated with diazepam (Fig. 3d) as determined by a two-sample Kolmogorov-Smirnov test (*n* = 4, *P* < 1e-3). Interestingly, the total number of spikes per unit remained similar across all organoids (Supplementary Fig. 7).

**Figure 3.**
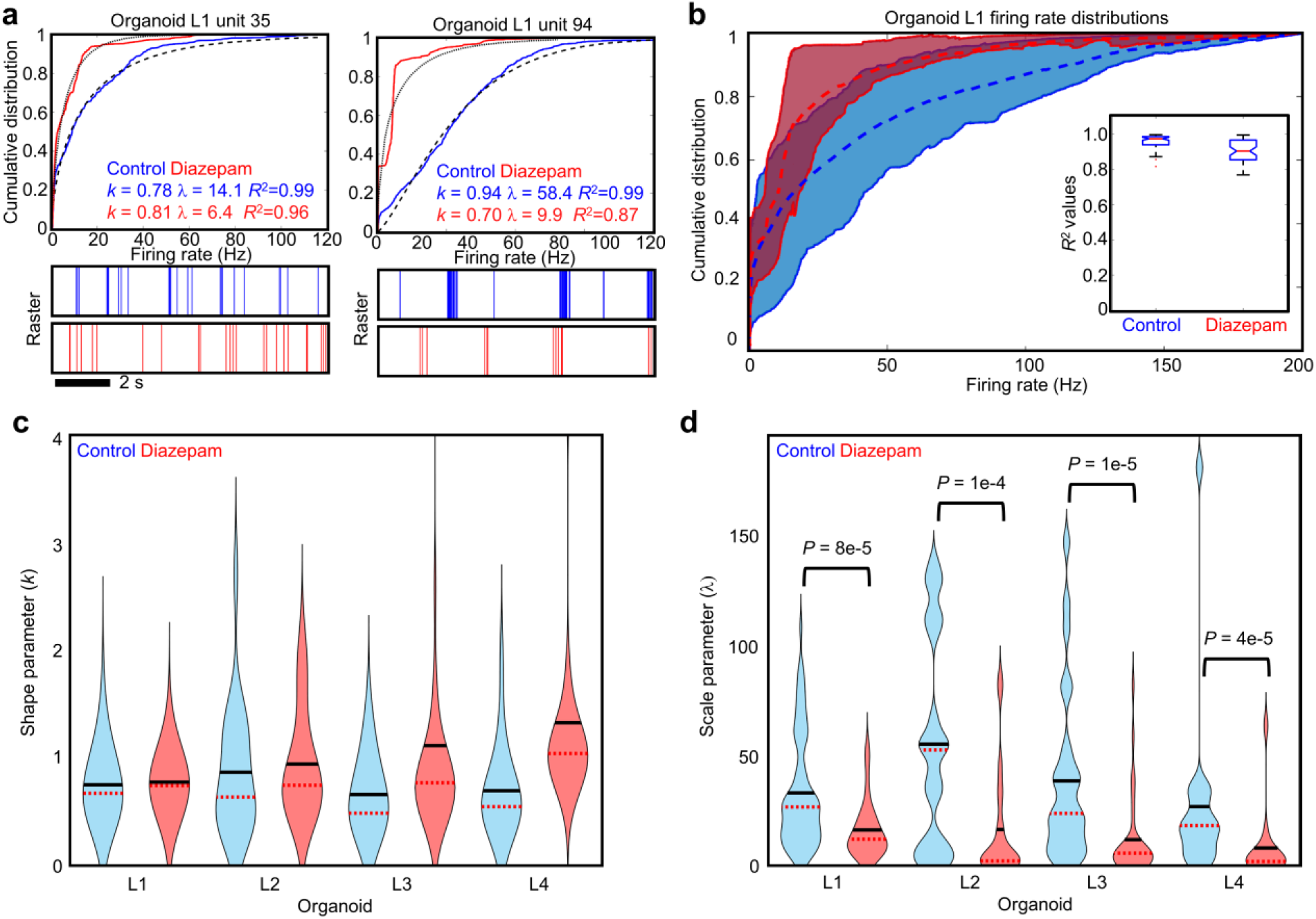
Neuron firing rates exhibit a characteristic non-linear response to a benzodiazipine. **a** Top, neuron firing rate cumulative distributions are shown for two representative neurons under control conditions (blue line) and after bath application of 50 μM diazepam (red line). The dotted lines are fits to a Weibull cumulative distribution function (CDF), where shape parameter (*k*) and the scale parameter (*λ*) are the only adjustable free fitting parameters. Raster plots are shown below for control (blue) and diazepam (red) conditions highlighting a 10 s window of spiking activity over the three-minute recording. **b** Cumulative firing rate distribution bounds are shown for organoid L1 under control (blue) and 50 μM diazepam (red) conditions. The shaded regions represent 80^th^ and 20^th^ percentile bounds across the spiking units, while the dotted lines represent the mean value. The inset shows the *R*-squared values of the Weibull CDF fits to the cumulative firing rate distributions for control and diazepam conditions highlighted in **a**. **c** The shape parameter *k* remains similar across organoid samples. The majority of the distributions remain below one, although organoid L4 does show a slightly larger increase in *k* with diazepam relative the other organoids. **d** The firing rate distribution scale parameter l is however significantly attenuated (visualized as violin plots), as determined by a two-sample Kolmogorov-Smirnov test (*P* < 1e-3), relative to control conditions for *n* = 4 organoids.

Concomitant changes were observed in single-unit spiking activity at the population level (Fig. 4). Temporal dynamics of coordinated neuronal bursting activity are visualized by the population averaged firing rate (averaged over a 5 ms Gaussian kernel, see Methods) for all bursts relative to their peak onset times (Fig. 4a, gray lines). Burst onsets initiate rapidly (within 100 ms) and persist for several hundreds of milliseconds with variable activity before subsiding. The variation in burst-to-burst activity is visualised by the standard error of the mean (SEM) across all burst events (Figure. 4a, red line). For control conditions the SEM peaks during the burst onset and reaches a steady state value during the duration of the burst. However, under diazepam (50 μM) the burst-to-burst SEM remains, on average, at a lower value. Figure 4b highlights the average difference in population averaged firing rates (averaged over the blue dotted intervals in Fig. 4a) across all burst events occurring during control and 50 μM diazepam treatment. Interestingly, diazepam reduced variation in the temporal structure of population-level activity across organoids (Fig. 4c) when averaging over all burst epochs.

**Figure 4.**
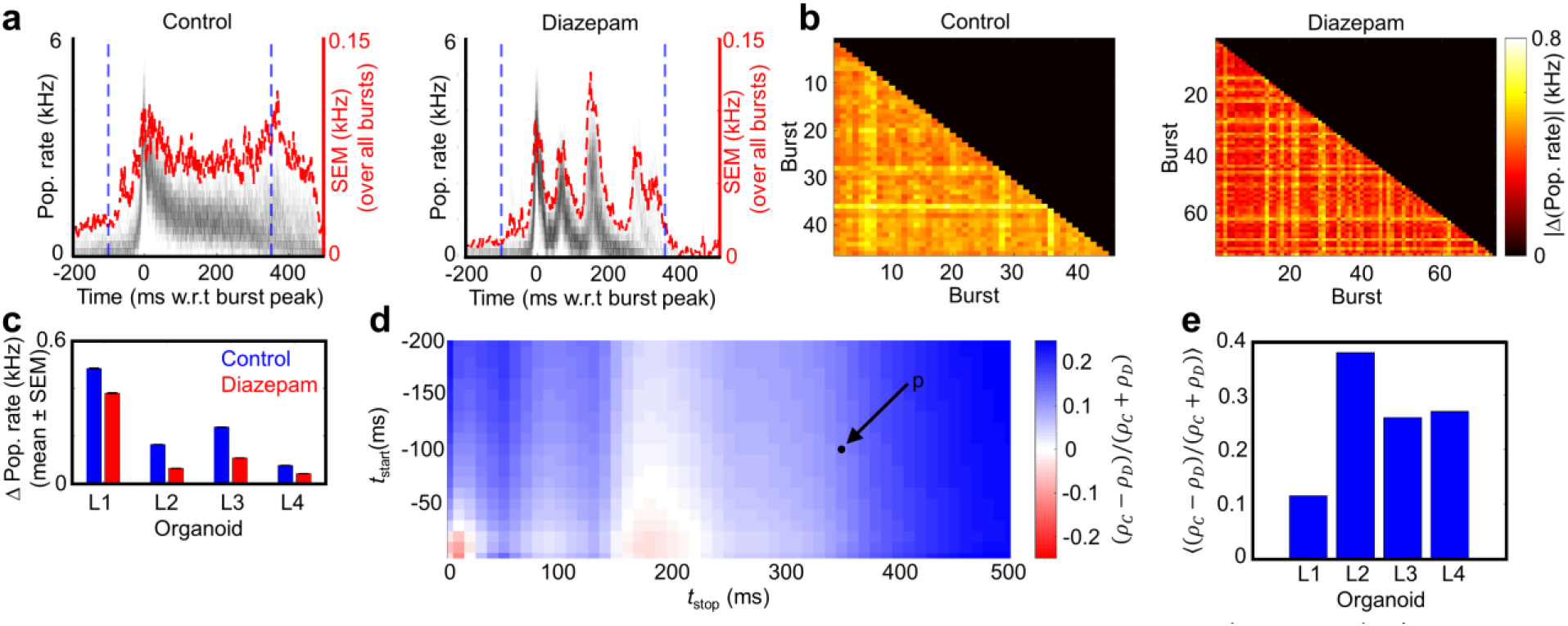
Diazepam-induced changes in population-level dynamics. **a** The population averaged firing rate (pop. rate) for the control and diazepam recordings (organoid L1 here) calculated from single-unit activity averaged over a 5 ms window. The population rate for each burst is plotted individually, centered by the peak in multi-unit activity (MUA). The standard error of the mean (SEM) calculated over all bursts is plotted in red. **b** Population rate average differences are shown across all individual burst pairs for the control and diazepam (50 μM) recordings from the same organoid. The average population rate difference is computed over a time window of −100 ms to +350 ms relative to the MUA peak (blue dotted line in **a**). The color bar scale is the same for control and diazepam. See **Supplementary Fig. 9** for visualization from a separate organoid. **c** Average population vector differences taken over all individual burst pairs per recording. The population vector difference is computed over a time window of −100 ms to +350 ms relative to the MUA peak (blue dotted line in **A**). Error bars indicate the SEM over all individual burst pairs. **d** Fractional change between population rate for control and diazepam (*ρ_C_* – *ρ_D_*) /(*ρ_C_* + *ρ_D_*), averaged over a range of different time windows for L1. Here, *ρ_C_* and *ρ_C_* are the average population rate differences for control and diazepam, respectively. Blue tiles indicate a higher average population vector difference for control. The black dot (*P* = (−100, 350 ms)) indicates the time window used in **b** and **c**. **e** Average score of the matrix in **d** for each organoid.

To further dissect variation in burst-to-burst activity we examined the fractional change in the average population rates over a given time interval relative to the multi-unit activity (MUA) burst peak (Fig. 4d), given by the following expression (*ρ_C_* – *ρ_D_*) /(*ρ_C_* + *ρ_D_*). Here, *ρ_C_* and *ρ_D_* are the average population rate differences for control and 50 μM diazepam conditions, respectively, for the given time interval relative to the burst peak set by *t*_start_ and *t*_stop_. Overall, we observe increased burst-to-burst variability when comparing control conditions relative to diazepam (across the full burst window); however, there is a slight increase in diazepam variability for organoid L1 over the narrow interval [150, 200] ms relative to the burst peak (this increase was not present in other organoids, see for example Supplementary Fig. 9). When averaging over all burst intervals (all matrix elements shown in Fig. 4d) we observed a consistent increase in burst-burst variability across organoids for control conditions relative to diazepam (50 μM) treatment (Fig. 4e). These observations, combined with significant decrease in distribution of single-unit firing rates (Fig. 3d), suggested that diazepam skews neuronal activity in our organoids toward a more redundant and strongly interconnected network with the capacity to instantiate fewer states defined as active functional relationships.

### Short-term interactions in human brain organoids

Neurons found in cortical circuits form local networks that exhibit robust short-term interactions with one another as quantified by correlations in spiking activity ^37–39^ and are likely to be mediated synaptic transmission ^40–42^. To determine if similar interactions occurred among neurons in our organoids, we constructed a map of spike correlations between single-unit spike trains and determined the latency distribution between correlated single-unit spike trains (Figure 5a,b). A prominent feature is that the distribution of pairwise mean spike latencies are short (4.7 ± 2.3 ms (mean ± STD) measured across 6 organoids from 14 separate measurements), as observed in local neocortical circuits *in vivo* ^37^. The correlation magnitude between two pairwise spike trains was computed using the spike time tiling coefficient (STTC) (Fig. 5c), which takes into account neuron firing rates, that can confound traditional cross-correlogram approaches ^38^ and generates weighted pairwise spike correlations (Figure. 5d). The edge weights were determined from the STCC (Fig. 5d, line thickness) using a correlation window size of *Δt* = 20 ms (the correlation values reach a stable plateau for *Δt* ≥ 20 ms (Fig. 5c), consistent with the histogram in Fig. 5b). The relative delay times between putative connections identified by correlated spike trains and spike-time jitter resulted in a mean latency within ≈ 5 ms time window with very few latencies longer than 10 ms (see Supplementary Fig. 10a for latency distributions from multiple organoids). These results suggested functional coupling between neurons within the organoid and were consistent with the unidirectional nature and timescales of synaptic transmission ^37,39, 43–45^.

**Figure 5.**
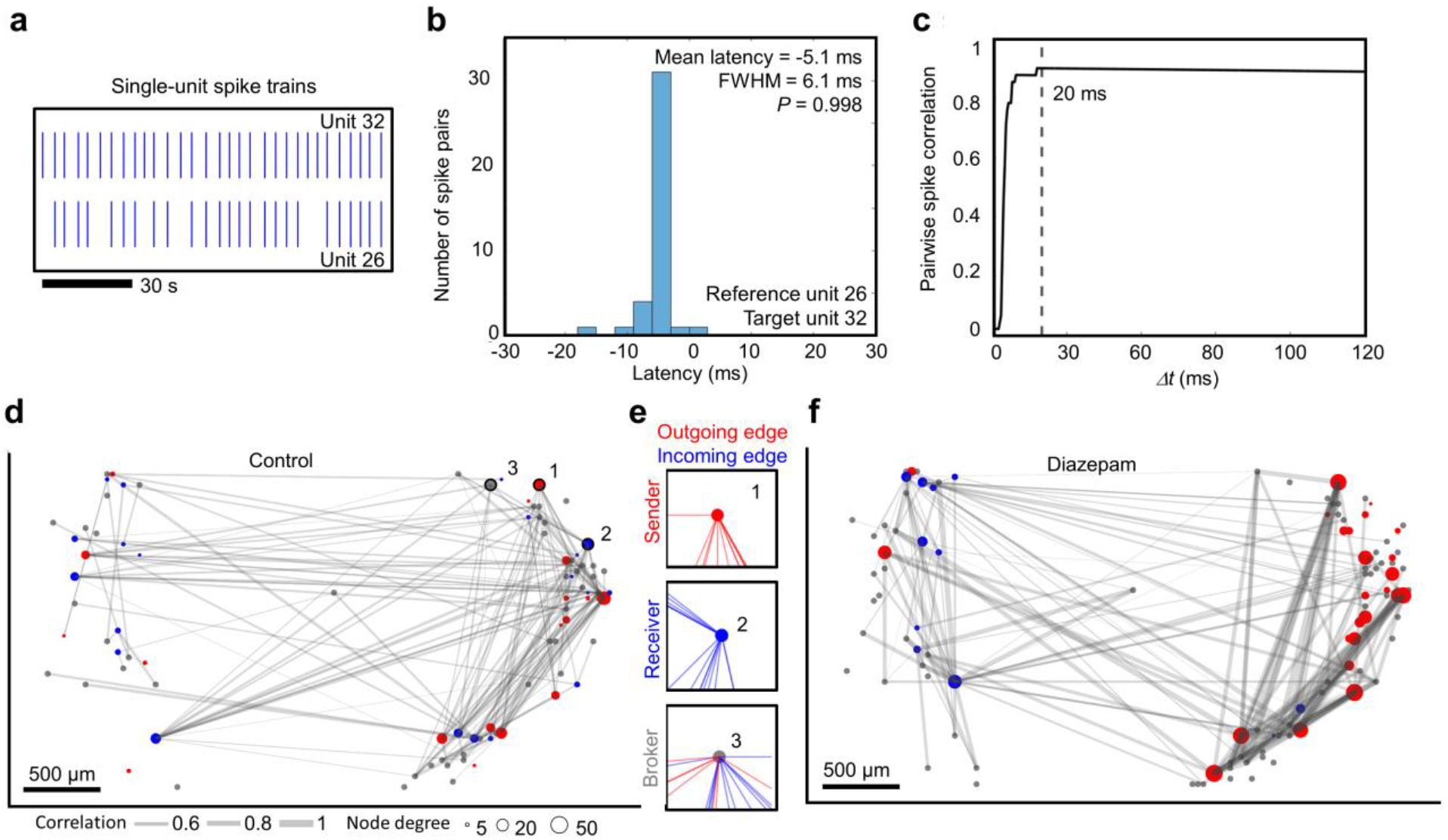
Mapping information flow through a human cerebral organoid. **a** Extracellular action potential spike events are shown from two correlated spike-sorted units. **b** The spike time latency distribution is shown between the two spike-sorted units shown in **a**. The latency distribution between pairwise spike events is unimodal (Hartigans’ dip test for multimodality, *P* = 0.998), with a mean latency of 5.1 ms and a FWHM of 6.1 ms. **c** The pairwise spike correlation was calculated using the spike time tile coefficient (STTC) as a function of correlation time window (Δ*t*) for the two correlated spike-sorted units shown in **a**. Choosing a correlation time window Δ*t* =20 ms captures all pairwise spike interactions between the two units. **d** Functional connectivity map showing the pairwise correlation strength (edge thickness in gray) between spike trains. Sorted by directionality, the in-degree and out-degree were computed per unit, defined as predominately incoming or outgoing edges respectively and designated *receiver* (blue) nodes, *sender* (red) nodes. All other nodes were labelled *brokers* (gray with a fixed size not indicative of node degree). For visual clarity, only the top 90 outgoing and the top 90 incoming edges are shown for sender and *receiver* nodes, respectively. **e** Examples of single *sender* (1), *receiver* (2) and *broker* (3) nodes showing all incoming (blue) and outgoing (red) edges for the spatial sites identified on panel **d**. The relative fraction of sender, receiver and broker edges remained similar across multiple organoids (*n* = 4, **Supplementary Fig. 11a**). **f** Functional connectivity map of the same organoid after treatment with 50 μM diazepam.

Predominant directionality was assigned to the edges of the functional connectivity map based on the spike latency distributions between single-unit pairs (see Methods for additional details). A negative mean spike time latency compared to the reference unit indicated a predominant directionality towards the reference unit and vice versa (Fig. 5b). Bidirectional connections as suggested by non-unimodal latency distributions were excluded from the analysis and constituted a small fraction of the total identified connections (7.13% ± 5.06% *n* = 14 measurements across *n* = 6 organoids). Based on the directionality, the in-degree (*D*_in_) and out-degree (*D*_out_) were computed per unit, defined as the number of incoming and outgoing edges respectively. Nodes with a high fraction of outgoing edges (*D*_out_ – *D*_in_)/ (*D*_out_ + *D*_in_) > 0.8 were labelled ‘s*ender*’ nodes (Fig. 4e, top). Nodes with a high fraction of incoming edges (*D*_in_ – *D*_out_)/ (*D*_out_ + *D*_in_) > 0.8 were labelled ‘*receiver*’ nodes (Fig. 5e, middle). Differences in the directionality vector less than 0.8 were labelled ‘*broker*’ nodes (Fig. 5e, bottom), which represented the majority of the nodes (*senders*: 15.4% ± 2.6%, *brokers*: 62.7% ± 6.0%, r*eceivers*: 21.9% ± 6.5% (mean ± STD)). These data were computed from four independent organoids (Supplementary Fig. 11a).

When assembled as described above, the spike time correlation map revealed a traffic pattern through the organoid visualized as a directional graph (Fig. 5d). The largest component— a graph of contiguous interconnected nodes—was parameterized according to its connection strengths as determined from pairwise spike correlations between spike trains. Setting the lower limit of connectivity strength at 0.35 minimized spurious connections (see Methods for randomization details and Supplementary Fig. 10b), and the number of active units (nodes) and edge (connection) strengths distribution were stable (> 4 hours) in that they did not vary significantly under control conditions (two-sample KS-test; *P* ≥ 0.1) (Supplementary Fig. 12). Edge strengths did not fit a random distribution (Fig. 6a, Supplementary Fig. 13). Rather, a truncated power law ^46^ and a gamma distribution fit experimental data well, with the gamma distribution outperforming the truncated power law (Methods and Supplementary Fig. 13). This result was consistent with observations that describe connectivity in cortical networks as a “skeleton of strong connections among a sea of weaker ones” ^42,47^.

**Figure 6.**
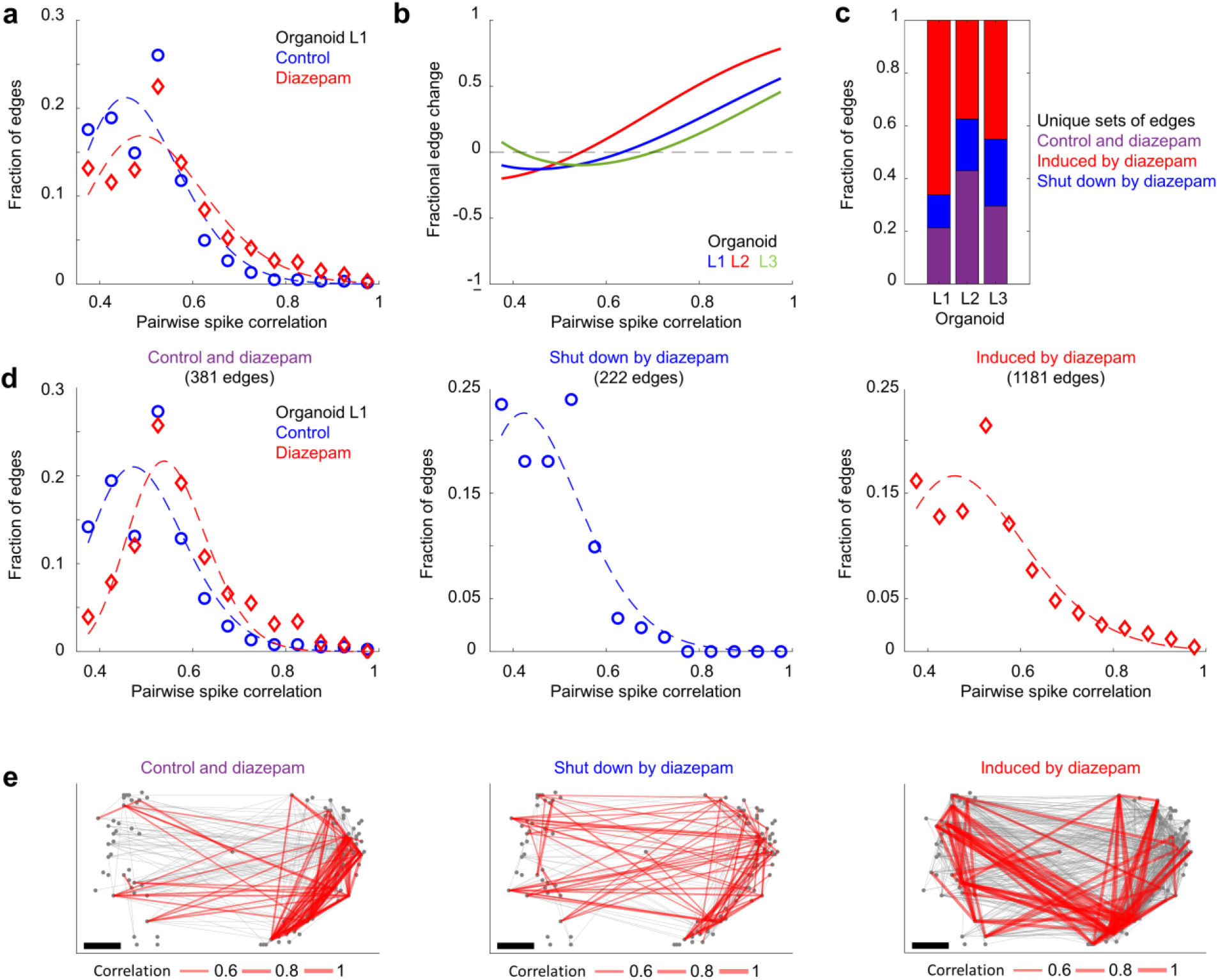
Diazepam selectively modulates the skeleton network of strong connections. **a** Pairwise spike correlation (edge) distributions of the functional connectivity maps shown in Fig. 5d,f. The edge strengths (of the largest component of interconnected nodes) were binned using a bin size = 0.05 over a connectivity range from 0.35 to one. A gamma distribution fit illustrates “a skeleton of stronger connections immersed in a sea of weaker ones”. Diazepam (50 μM) increases the minority population of stronger connectivity strengths, while decreasing the connection strengths of the weaker and more abundant connections with respect to control conditions. **b** The fractional difference in edge strengths, (*f*_d_ – *f*_c_)/(*f*_d_ + *f*_c_), is shown, where *f*_c_ and *f*_d_ are the gamma distribution fits highlighted in panel **a** for control and diazepam (50 μM) conditions, respectively. A higher proportion of strong connections are present in diazepam (50 μM) conditions relative to control conditions for *n* = 3 organoids. **c** The relative fractions of unique sets of edges are shown. Purple bars represent edges present during both control and diazepam conditions. Blue bars represent the fraction of edges shut down by diazepam and the red bars represent the fraction of edges induced by diazepam. **d** Edge strength distributions highlighted in **c** are plotted for organoid L1, while the spatial graph is shown in **e**. The top 90 edges (by weight) are shown in red, while the remaining edges are plotted in gray.

Diazepam (50 μM) strengthened the skeleton of strong connections while weakening the sea of weaker connections (Fig. 6a). The change in the fraction of edge strengths was further visualized as a fractional difference (*n* =3 organoids, Fig. 6b, organoid L4 was excluded due low number of edges relative to the other samples) defined as (*f*_d_ - *f*_c_)/(*f*_d_ + *f*_c_), where *f*_c_ and *f*_d_ are the fraction of total edges as a function of pairwise spike correlation for control and diazepam (50μM), respectively. Remarkably, we found that diazepam shut down a minority population of more weakly connected existing edges (Fig. 6c), while inducing a new, more strongly interconnected subnetwork (Fig. 6c-e). Together, these results combined with an increase in the density of functional connections under diazepam (50 μM) treatment (Supplementary Fig. 11b), reveal a transition to more correlated and interconnected network that exhibits less variable single-unit level (Fig. 3) and population-level neuronal dynamics (Fig. 4) relative to control conditions in the organoids. Of relevance is that diazepam can increase resting-state functional connectivity in the medial visual system, as measured by fMRI in healthy volunteers ^48^, which is a brain region rich in GABA_A_ receptors and shown to have high binding of GABAergic drugs.

### Theta frequency oscillations

To extract oscillatory activity from the extracellular field, we decomposed the broad-band voltage vs. time signals into local field potential (LFP) frequency sub-bands by band-pass filtering of raw data (Fig. 7a, see Methods). Although we observed LFPs at multiple frequencies (Supplementary Fig. 14), we limited this analysis to theta oscillations (4-8 Hz) that were considered significant over time intervals when the oscillation amplitude envelope exceeded the noise floor (Supplementary Fig. 15-16). Multiple brief theta cycles occurred at different spatial locations across the organoid (Fig. 7b, red lines) that can be referenced to burst activity (Fig.7b, black line) signified when the MUA population-averaged spike rate peaked within the burst. These theta oscillations showed consistent phase offsets relative to each other (Fig. 7c, gray lines). When we blocked AMPA, NMDA and GABA_A_ receptors along with TTX-sensitive sodium channels, the LFP amplitudes were no longer observable above the CMOS amplifier noise floor (*n* = 4 organoids; Supplementary Fig. 15). This observation validated the finding that LFP signals depended upon synaptic transmission-based network activity ^49^.

**Figure 7.**
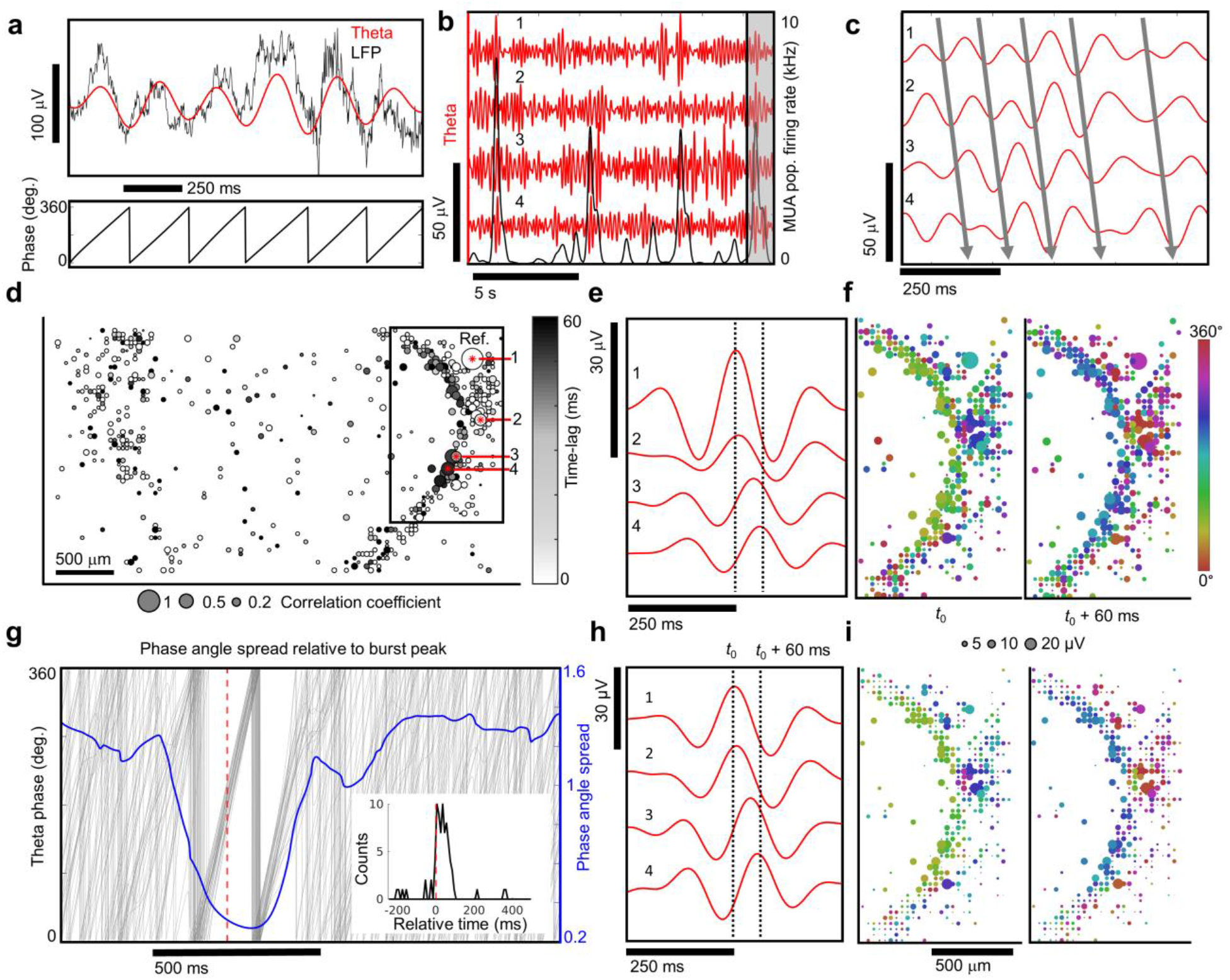
Spatial and temporal coherence of theta oscillations with neuronal population bursts. **a** The raw local field potential (< 500 Hz, black line) and the 4-8 Hz theta filtered band (red) (top). The phase of the theta oscillation (bottom). **b** Theta band oscillations (red) from four different recording sites and the multi-unit activity (MUA) population averaged firing rate (black) averaged over a 100 ms window. **c** Zoomed in view of highlighted black rectangle in **b**. The solid lines indicate the relative phase offsets of theta oscillations across spatial sites of the organoid. Within a narrow time window these oscillations showed consistent phase offsets. **d** Spatial correlation map of theta oscillations. The correlation coefficient (bubble size) is shown with respect to the seed reference site (1) and the relative phase-lag with respect to the reference electrode shown in grayscale reveals spatial alignment of theta oscillations. **e** Signal averaged theta oscillations using peaks from electrode (1) as a reference. The numbers 1-4 in **c-e, h** all refer to the same set of electrodes. **f** Spatial map of signal averaged theta oscillation phase and amplitude relative to reference electrode number 1. Two time points are shown, one at the center of the reference electrode *t*_0_ and another 60 ms later. **g** Phase angle spread relative to burst peak (blue line). Individual theta phase traces (gray line) are plotted for electrode 1 relative to population burst events determined from MUA averaged over a 5 ms window. The phase angle spread is minimized after the burst peak (red dotted line). The time of the theta peak amplitude relative to bursts where the angular spread is minimized across multiple electrode sites (inset). **h** The same theta oscillations from **e** are signal averaged with respect to population burst peak times (*t*_0_). **i** Spatial map of signal averaged theta oscillations using population burst peak times reveal a temporal alignment of theta oscillations with neuronal population bursts. Panels **f**, **i** share the same scales.

Cross-correlation coefficients of theta oscillations and their relative time-lags between all electrode pairs were used to generate a spatial correlation map with a focal region of highest correlated activity and their phase delays relative to more distant positions on the map (Fig. 7d). Signal averaging theta oscillations with respect to oscillation peaks above the noise floor from the highest correlated node (identified in Fig. 7d) further validated synchronized theta activity with relative phase shifts across the organoid (Fig. 7e,f see Supplementary Fig. 17 for visualizations in a different organoid). We further demonstrated that LFPs were synchronized with neuronal population bursts (see next section). We confirmed the validity of the LFP theta signal by estimating imaginary coherence, a metric, which is dependent on non-zero phase synchrony ^50^, and found sets of highly coherent electrodes (Supplementary Fig. 18), which corresponded to the same spatial regions identified by cross-correlation and signal averaged analysis (Fig. 7d-i). Diazepam induced a dramatic reduction in pairwise theta correlations (Supplementary Fig. 19d) and a significant decrease in imaginary coherence across the organoid (*n* = 3 organoids, paired t-test, *P* < 1e-4, Supplementary Fig. 18f).

### Peak theta amplitudes are synchronized with neuronal population bursts

Spatiotemporal synchronization of local neuronal populations generate oscillatory rhythms observed in the LFP ^49^. The amplitude of these oscillations are correlated with an increase in local population-level spiking activity, as observed *in vivo* ^51,52^. We also observed a sharp increase in the amplitude of theta oscillations occurring during time periods exhibiting synchronized neuronal population bursts (Supplementary Fig. 16). First, we identified the time points when the neuronal population firing rates (MUA averaged over a 5 ms time window) peaked within burst epochs. Burst-peak times were then used as temporal anchor points to signal average theta oscillations (Fig. 7g,h) across all electrode sites. A spatial map (Fig. 7i) of signal-averaged theta oscillations (relative to the MUA population burst-peak) revealed regions of theta activity strongly correlated with burst peaks. Signal averaging theta amplitudes relative to population burst-peaks revealed the same spatial pattern of theta phase coherence attained by independently signal averaging theta oscillations relative to their peak oscillation amplitude time points (Fig. 7e,f). Theta phase alignment was maximized within 0-100 ms after to the population burst peak (red dotted line is the burst peak in Fig. 7g and inset) and shown by a sharp drop in the circular standard deviation of the phase (phase angle spread). Temporal windows of constant phase shifts (defined where the circular standard deviation of the theta phase dropped below one) persisted over a ≈ 400 ms time window (roughly 2-3 theta cycles). The time spent above the noise floor substantially increased during broader burst periods (*n* = 4 organoids, Supplementary Fig. 16). Modulating brain organoid signaling with diazepam (50 μM) induced a notable reduction (>30%) in the number of correlated theta electrode sites (above a correlation threshold of 0.2) across the organoids (*n* = 3) (Supplementary Fig. 19d), a result consistent with the known down-regulation of theta activity by benzodiazepines *in vivo* ^53,54^. Taken together, these results show that theta oscillation amplitudes are correlated with neuronal population bursts.

### Spike phase-locking to theta

In the absence of sensory input brain activity can entrain action potentials to theta frequency oscillations ^55^ and define a temporal window for spiking within local circuits ^56,57^. Previous work *in vivo* and *ex vivo* has relied on manually positioned, low-density recording electrodes, to identify a handful of units exhibiting preferential spike phase-locking to theta frequencies at a given moment in time ^55,56,58,59^. We looked for relationships between theta frequency oscillations and single-unit spikes in organoids using the high density and large spatial extent of the CMOS arrays. To determine whether spikes occurred at a preferred theta phase, we counted spikes in relation to the phase of theta cycles (across 1,020 electrodes) (Fig. 8a) and determined significance with the Rayleigh criterion for non-uniform distribution of spikes in circular phase space ^55^. By selecting the top 1,020 microelectrodes (out of 26,400 based on spike activity), we simultaneously surveyed both theta oscillations and spikes across the organoid to draw a spatial map of the mean phase-locked spike angle to theta for regions of highly correlated theta activity identified previously by the cross-correlation analysis (Fig. 8c and Supplementary Fig. 20 for analysis from *n* = 3 organoids). Theta-phase spike event distributions for electrode locations revealed a strong preference for spiking during a narrow phase window of theta centered about its mean angle μ (Fig. 8b). Theta oscillations can provide a temporal window for computation within local circuits *in situ* ^56,60^ suggesting that organoids may display sufficiently complex circuitry for intrinsic computation.

**Figure 8.**
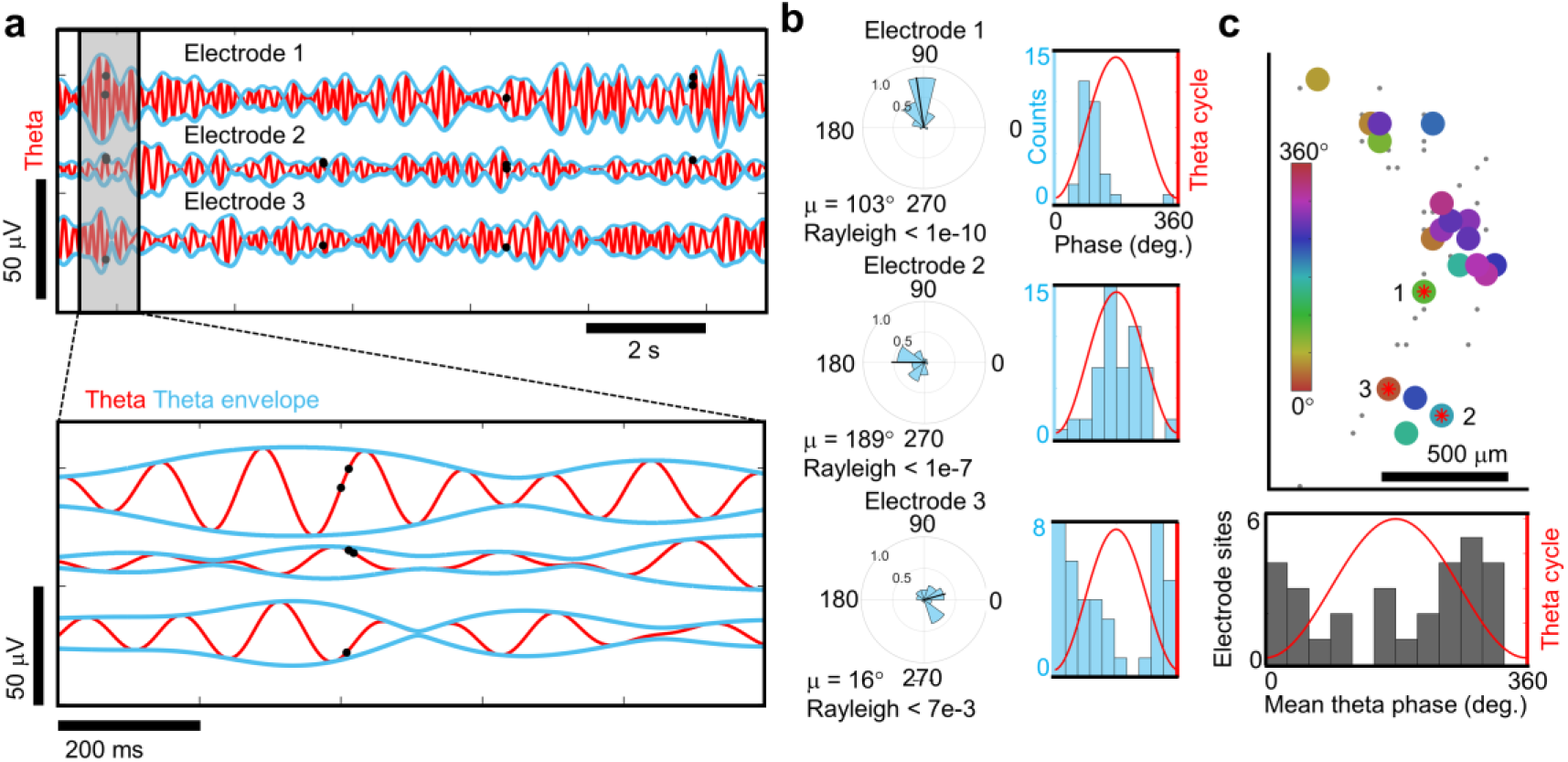
Phase-locking of spikes to theta oscillations. **a** Top, theta oscillations (red line) from three representative electrode sites and the single-unit spikes that occurred at each of those electrodes (black dots). Bottom, zoomed in view from the gray boxes on the top. **b** Left, circular distribution of theta phase angles occurring during single-unit spike events measured from the electrodes in **a**. Direction of the mean spike angle (μ) relative to the theta phase and magnitude (mean resultant length) are shown in the polar plots. The Rayleigh criteria for non-uniformity was used to determine if spikes were distributed non-uniformly over the theta cycle (0°, 360°). Right, distributions of theta-spike angles shown relative to the theta cycle are visualized as histogram plots. **c** Top, a cluster of phase-locked units to theta oscillations are shown within the coherent pocket highlighted by the box in Fig. 7d. The color indicates the mean phase-locked angle to theta (μ) that satisfy the Rayleigh criteria for non-uniformity (*P* < 0.05). Electrode sites with no preferred theta phase (*P* > 0.05) are shown by gray dots. Bottom, histogram of the mean phased-locked angle to theta for all electrode sites across the array that satisfy the Rayleigh criteria (*P* < 0.05).

### Acute measurements in whole organoids

To determine that spiking activity extended in the z-plane of the organoids, as opposed to the 2D recordings from slices on 2D CMOS arrays, we performed acute extracellular recordings from three intact brain organoids with a densely tiled 960 electrode CMOS shank with ≈20 μm inter-electrode pitch (Neuropixels probe; Jun et al., 2017). Recordings were made by inserting the CMOS shank into the organoid using a motorized, low-drift micromanipulator (see Methods). We observed single-unit spiking activity that extended hundreds of microns across the organoid tissue within a limited stratum of the z-plane in three whole organoids (Fig. 9a,b). Similar to the results from organoid slices, we observed spontaneous, synchronized population bursts.

**Figure 9.**
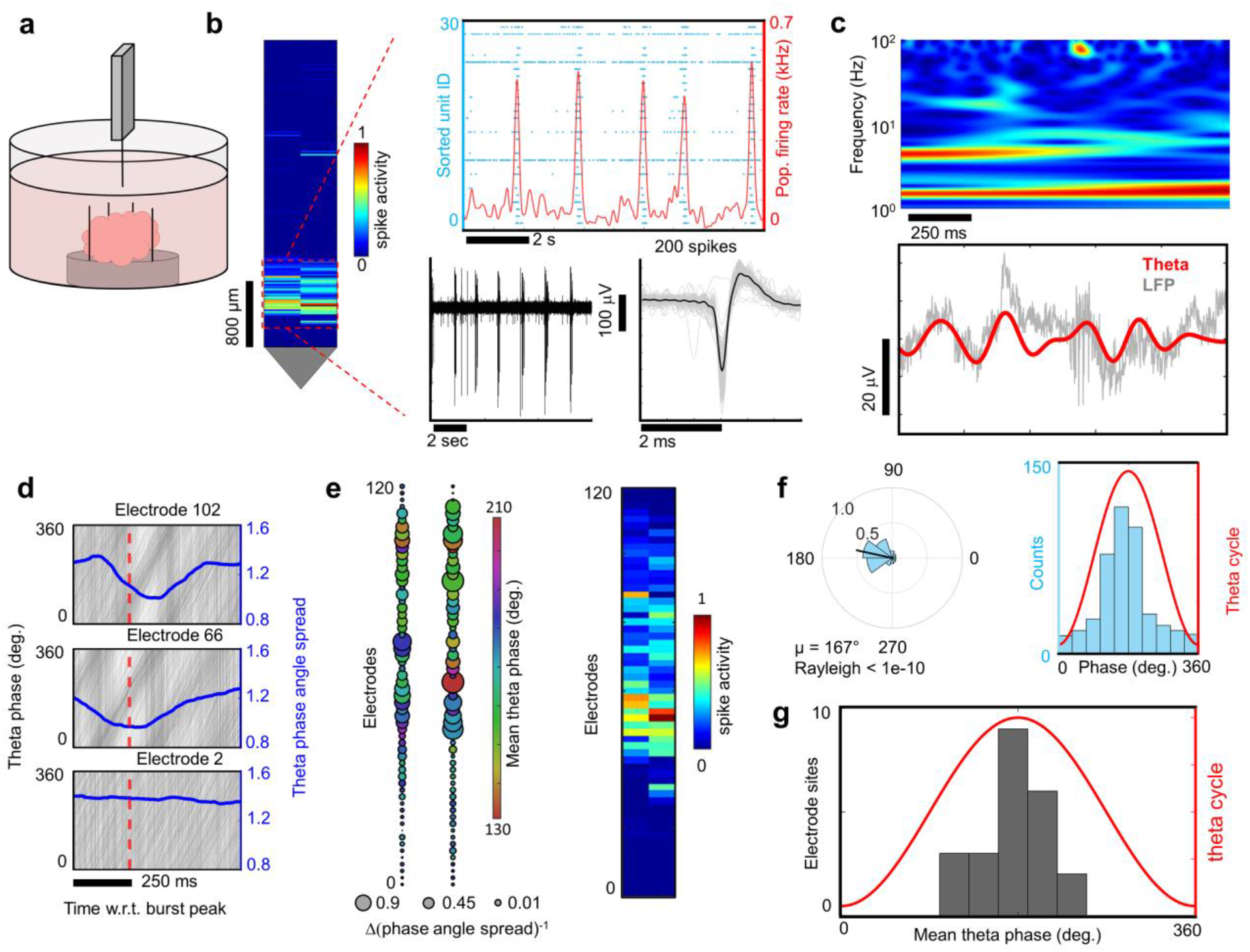
Neuropixels CMOS shanks resolves spiking and LFP in the z-plane. **a** A Neuropixels high-density CMOS shank was attached to a custom-made mount and controlled by a micromanipulator in order to lower the shank into an immobilized brain organoid kept at 37 °C in BrainPhys media. **b** Left, spikes (above a 5-rms threshold) were detected in a subset of electrodes near the tip of the shank. Spike activity is normalized relative to the electrode with the most detected spikes. Top right, raster-plot visualization of single-unit spiking activity (Kilosort2) as shown by the blue dots for each sorted unit. Stereotyped population bursts are visualized as peaks in the population averaged firing rate (red line). Bottom right shows the extracellular field potentials (0.3 - 4 kHz) generated by a spiking neuron within the organoid as measured by the shank. **c** Top, spectrogram plot of local field potential (LFP) from an electrode illustrates dominant oscillation power in the theta and delta bands. Bottom, raw LFP (gray line) from the same electrode overlaid with the theta-filtered band (red line). **d** Theta phase traces (gray lines) are shown relative to population burst peak events (red dotted line) reveal phase coherence as illustrated by a drop in the phase angle spread (blue line). The bottom plot shows an electrode site displaying no phase coherence relative to population burst events. **e** Left, spatial map of the change in theta phase angle spread is shown relative to burst peak time across the shank. The bubble size indicates the inverse of the phase angle spread relative to the burst peak for each electrode. Notice the overlap with phase coherent sites and the spatial region registering spiking activity on the right. **f** Left, circular distribution of theta-spike phase angles measured from a single electrode site. The direction of the mean spike angle (μ) relative to the theta phase and magnitude (mean resultant length) are shown as polar plots. The Rayleigh criteria for non-uniformity was used to determine if spikes were distributed non-uniformly over the theta cycle (0°, 360°). Right, distributions of theta-spike angles shown relative to the theta cycle. **g** Mean theta phase angle of spike phase-locked electrodes across the Neuropixels probe satisfying the Rayleigh criterion (*P* < 0.05).

We further utilized the Neurpixels shank to confirm synchronization of theta oscillations to neuronal population bursts and identified electrode sites along the z-plane that exhibited prominent theta band oscillations in the LFP (Fig. 9c). To determine whether the population bursts were phase-synchronized to theta oscillations in the intact organoids, we signal-averaged theta phases relative to MUA population burst peak times along the shank (Fig. 9d,e). Theta phase alignment was synchronized with population bursts and co-localized with spatial regions where spiking activity was observed (Fig. 9e, right). Meanwhile theta phases added deconstructively in regions where the organoid showed no signs of spiking activity. Our measurements of electrical activity generated by brain organoids provided strong evidence that theta is synchronized over temporal epochs of increased neuronal activity within the time window of population bursts ^49^. Finally, utilizing acute field recordings (with sufficient bandwidth to simultaneously resolve both action potentials and local field potentials) from whole brain organoids, we confirmed phase-locking of spikes to theta (satisfying the Rayleigh criterion) (Fig. 9f) simultaneously across multiple sites (Fig. 9g) in the z-plane of organoids. The acute high-density extracellular field recordings, described above, captured a window of neuronal activity in the z-plane using a probe with a small cross-sectional footprint (70 x 24 μm). The presence of both spiking and LFP activity were observed verifying that network activity occurred within a limited stratum several hundred microns within the 3D structure of the organoid and reflect the activity observed on planar high-density arrays.

## Discussion

Human brain organoids spontaneously self-organize their physiologic activity such that significant non-random coupling occurs between single-units. Over the entire area of the organoid assessed by the MEA this activity has the form of a network in which single-units are considered nodes connected by weighted directionally defined edges ^37,39,42^. The robust detection of signals was possible with high-density CMOS microelectrode arrays that record spiking with high temporal and spatial resolution over a sufficiently large section of tissue to capture theta oscillation coherence and phase-locking to spikes. When cultured for prolonged time periods organoids can undergo some degree of neuronal maturation, including the elaboration of dendritic spines and the formation of spontaneously active neuronal networks ^5^. Indeed, our single-cell analysis and immunohistochemistry demonstrated cellular diversity that is collectively capable of supporting LFPs (Fig. 2 and Supplementary Fig. 5,6). For example, a variety of inhibitory neurons were present, including parvalbumin cells (Fig. 2c) which are associated with theta oscillations ^61^ and axon tracts that extended over millimeters (Fig. 2b). Single-cell transcriptomics indicate the presence of functioning cells capable of forming complex neural networks including the presence of GABAergic cells that are consistent with their role in the generation of highly correlated activity networks detected as LFPs ^27^.

The formation of a connectivity map (Fig. 5d) demonstrated that the organoid was capable of establishing a weighted network of inflow and outflow tracts over short and long distances. As demonstrated in the murine brain ^42,47^, high connection strength edges shape a non-random framework against a background of weaker ones (Fig. 6a, Supplementary Fig. 13). The majority of the nodes, which we refer to as brokers, have large proportions of both incoming and outgoing edges. The dynamic balance among receivers and senders could calibrate a global mechanism for setting the brain state of activation.

We developed quantitative metrics that capture changes in single neuron firing patterns, networked correlations between neuron pairs and aggregate population-level dynamics simultaneously across millimeter spatial scales. Furthermore, these cerebral organoids— composed of roughly one million cells—have neuronal assemblies of sufficient size, cellular orientation, connectivity and co-activation capable of generating field potentials in the extracellular space from their collective transmembrane currents. Because the measurements were conducted within a small tissue volume (**≈** 3.5 mm^3^) the problem of volume conduction from distant sources, as happens in MEG measurements, is minimal. Correlations between theta oscillations and local neuronal firing strongly supported a local source for the rhythmic activity^49^. Consistent with minimal volume conduction effects, we validated theta oscillations by demonstrating that the imaginary part of coherency ^50^ explained the entirety of the coherency. The local volume through which theta dispersed extended to the z-dimension as shown with the Neuropixels shank (Fig. 9). Finally, we show that subpopulation of neurons within organoids exhibit phase-locking to theta oscillations (Fig. 8). These recordings span multiple spatial and temporal scales and so go beyond previous studies on organoid electrophysiology ^5,11,14,16^.

Spontaneous cortical activity can occur in reduced preparations such as slice cultures and acute slices; however, in such settings the brain tissue has had the experience of an input from the outside world and an output to a body. The brain organoid presents the unique circumstance of brain development devoid of any externality. Therefore, spontaneous activity in the organoid is *ab initio*, in contrast to the spontaneous activity arising in the cerebral cortex as the “resting state.” A neuronal system in which activity occurs in the absence of sensation fits what O’Keefe and Nadel reference as a physiologic scaffold capable of representing information from the external environment before it is experienced ^62^. Buzsáki makes a similar point ^63^ “…the brain already starts out as a nonsensical dictionary. It comes with evolutionarily preserved, preconfigured internal syntactical rules that can generate a huge repertoire of neuronal patterns.” However, Buzsáki makes the argument that meaning comes from motor activity. Regardless, the organoid, of course, is devoid of both sensory and motor input/output. The seeds of this concept existed long before organoids or even before neuroscience in the thinking of Immanuel Kant who described synthetic a priori knowledge as independent of experience, but hardly suspected a priori knowledge consisted of nothing more than abstract waveforms.

## Acknowledgments

The authors acknowledge the use of the William K. Bowes Laboratory for Stem Cell Biology and Engineering at the University of California, Santa Barbara. Illumina sequencing was carried out at the Biological Nanostructures Laboratory, within the California NanoSystems Institute, supported by the University of California, Santa Barbara and the University of California, Office of the President.

## Author contributions

T.S. and K.S.K. designed, conceived and supervised the study; P.K.H., K.R.T. and L.R.P. offered numerous suggestions and comments; T.S. designed and performed electrophysiology experiments on sliced organoids; T.S., T.V.D.M. and E.G. performed computational analysis and statistics on electrophysiology recordings; E.G. performed shank recording on whole organoids; S.M.K.G. cultured the organoids, performed single-cell RNA sequencing and analysis; A.P.B. performed spike sorting analysis under the supervision of A.H.; G.L. performed immunohistochemistry, imaging and organoid slicing; K.G.R. and K.K. performed imaginary coherence analysis under the supervision of S.S.N.; Z.C. performed additional analyses.; M.A. contributed to cell culturing; T.S. and K.S.K wrote the manuscript, and all authors discussed the results and commented on the manuscript.

## Data and materials availability

All data needed to evaluate the conclusions in the paper are present in the paper and/or the Supplementary Materials. Additional data and code related to this paper are available from the corresponding author upon reasonable request.

## Materials and Methods

### Cerebral organoid generation

The control iPCS line F12442.4 ^64^ was cultured in mTeSR1 medium (Stem Cell Technologies) on tissue culture plates coated with hESC-qualified Matrigel (Corning). mTeSR1 was exchanged every other day and iPSCs were routinely passed using ReLeSR (Stem Cell Technologies). Cerebral organoids were generated after the method by Lancaster *et al.* ^8^ with minor modifications. iPSCs were incubated in 0.5 mM EDTA in D-PBS for 3 minutes before dissociation in Accutase for 3 minutes at 37 °C. After adding mTeSR1 and triturating to achieve a single-cell suspension, cells were pelleted for 3 minutes at 1,000 rpm. Cells were resuspended in low bFGF hES media (20% KOSR, 3% ES-FBS, 1x GlutaMAX, 1x MEM-NEAA, 1x β-Mercaptoethanol, 4 ng/ml bFGF in DMEM/F12) supplemented with ROCK inhibitor (50 µM) and plated in U-bottom ultra-low attachment plates at 4,500 cells per well. On day 2, low bFGS hES media with ROCK inhibitor was replaced. Another media change was performed on day 4, with omission of bFGF and Rock inhibitor. On day 5, embryoid bodies were transferred to neural induction media (1x N2 supplement, 1x GlutaMAX, 1x MEM-NEAA, 1 μg/ml Heparin in DMEM/F12), and media was replaced on days 7 and 9. On day 10, each neuroepithelial structure was embedded in 15 μl hESC-qualified Matrigel followed by incubation in neural induction media for another 2 days. On day 12, neural induction media was replaced with NeuroDMEM-A media (0.5x N2 supplement, 1x B27supplement without Vitamin A, 1x β-Mercaptoethanol, 1x GlutaMAX, 0.5x MEM-NEAA, 250 μl/l insulin solution, 1x Pen/Strep in 50% DMEM/F12 and 50% Neurobasal). From day 19 on, organoids were cultured in NeuroDMEM+A media (0.5x N2 supplement, 1x B27 supplement with Vitamin A, 1x β-Mercaptoethanol, 1x GlutaMAX, 0.5x MEM-NEAA, 250 μl/l insulin solution, 12.5 mM HEPES, 0.4 mM Vitamin C, 1x Pen/Strep in 50% DMEM/F12 and 50% Neurobasal) with media changes twice per week. From day 21 on, organoids were kept on an orbital shaker at 75 rpm. Mycoplasma testing was routinely performed on organoids and iPSCs using the MycoAlert Mycoplasma DetectionKit (Lonza).

### Immunohistochemistry and microscopy

Organoid samples were fixed in 4% paraformaldehyde in 0.1M sodium cacodylate buffer (pH:7.4) overnight at 4 °C. After rinsing, samples were embedded 10% agarose and sectioned (100 µm) using a vibratome (Leica, Lumberton, NJ). Primary antibodies used were anti-Parvalbumin (1:200; abcam, Cambridge, MA; ab11427), anti-GAD65 (1:200; GeneTex, Irvine, CA; GTX113192), anti-SMI312 (1:500; BioLegend, San Diego, CA; 801701), anti-GFAP (1:500; abcam, Cambridge, MA; ab53554), anti-connexin 43 (1:200; Millipore, Burlington, MA; MAB3067), anti-MAP2 (1:500; GeneTex, Irvine, CA; GTX82661), anti-synaptobrevin (1:500; Synaptic Systems, Goettingen, Germany; 104-211). Samples were viewed through a 20x oil immersion lens (N.A. 0.85) and imaged using an upright Olympus Fluoview 1000 laser scanning confocal microscope (Center Valley, PA) equipped with a motorized stage. High-resolution wide-field mosaics were produced as described elsewhere ^65^.

### Single-cell RNA sequencing of cerebral organoids

Individual cerebral organoids were subjected to dissociation using the Worthington Papain Dissociation System (Worthington). Solutions were prepared according to the manufacturer’s instructions. Organoids in NeuroDMEM+A (defined in Cerebral organoid generation) media were cut into pieces with a sterile scalpel blade and washed twice in PBS prior to dissociation. Incubation in papain/DNaseI solution was performed at 37 °C and 5% CO_2_ with occasional agitation. After 30 minutes of incubation, organoids were triturated 10 times with a fire-polished glass Pasteur pipette, followed by further incubation and trituration every 15 minutes until dissociation was complete. The total incubation time in papain/DNaseI solution was 90 minutes. After adding Inhibitor Solution and DNaseI in EBSS, the dissociated cells were pelleted by centrifugation for 3 minutes at 300 g. The cells were resuspended in 500 µl ice-cold PBS/0.01% BSA using a wide-bore plastic pipette tip and passed through a 35 µm cell strainer. An aliquot of the single-cell suspension was mixed with Trypan blue and analyzed for viability and cell concentration on a Countess II FL Automated Cell Counter (ThermoFisher). Viability was > 90% for all organoid samples.

### Drop-seq library preparation

Following dissociation, cells were diluted to 110 cells/µl in PBS/0.01% BSA. Drop-seq was performed as described in Macosko *et al*. ^66^. Briefly cells and barcoded beads (Chemgenes, 132 beads/ul in lysis buffer) were run on an aquapel-treated microfluidic drop-seq device (FlowJEM) for co-encapsulation in nanoliter-sized droplets. After droplet-breakage, reverse transcription and Exonuclease I treatment, beads were counted, and 2,500 beads were apportioned per PCR tube for cDNA amplification. The amplified cDNA libraries were purified using SPRI beads (Beckman Coulter) and quantified on a Fragment Analyzer (Agilent). Tagmentations were performed using the Illumina Nextera XT kit (Illumina), and the resulting libraries were purified in two consecutive rounds of SPRI beads-based size selection (0.6x beads to sample ratio followed by 1x beads to sample ratio). The size and concentration of the final libraries were measured on a Fragment Analyzer and a Qubit Fluorometer (ThermoFisher), respectively. Libraries were sequenced on an Illumina Nextseq500 instrument.

### Drop-seq data analysis

Counts matrices were generated using the Drop-seq tools package ^66^. Briefly, raw reads were converted to BAM files, cell barcodes and UMIs were extracted, and low-quality reads were removed. Adapter sequences and polyA tails were trimmed, and reads were converted to Fastq for STAR alignment (STAR version 2.6). Mapping to the human genome (hg19 build) was performed with default settings. Reads mapped to exons were kept and tagged with gene names, beads synthesis errors were corrected, and a digital gene expression matrix was extracted from the aligned library. We extracted data from twice as many cell barcodes as the number of cells targeted (NUM_CORE_BARCODES = 2x # targeted cells). Downstream analysis was performed using Seurat 3.0^67,68^ in R version 3.6.3. An individual Seurat object was generated for each sample, and objects were merged using the Seurat merge function. Cells with < 300 genes detected were filtered out, as were cells with > 10% mitochondrial gene content. Counts data were log-normalized using the default NormalizeData function and the default scale of 1e4. The top 3,000 variable genes were identified using the Seurat *FindVariableFeatures* function (*election.method* = “vst”, *nfeatures* = 3,000), followed by scaling and centering using the default *ScaleData* function. Principal Components Analysis was carried out on the scaled expression values of the 3,000 top variable genes, and the cells were clustered using the first 30 principal components (PCs) as input in the FindNeighbors function, and a resolution of 0.4 in the *FindClusters* function. Non-linear dimensionality reduction was performed by running UMAP on the first 30 PCs. Following clustering and dimensionality reduction, putative cell doublets were identified using DoubletFinder ^69^, assuming a doublet formation rate of 5% and calculating homotypic proportions based on clustering at resolution 0.4 as described above. A pk value of 0.61 was selected for doublet identification based on the results of *paramSweep*_vs, *summarizeSweep* and *find.pK* functions of the *DoubletFinder* package. After filtering out the doublets as identified by *DoubletFinder*, counts data were extracted, and a new Seurat object was generated. Normalization, variable gene selection, scaling, clustering and UMAP dimensionality reduction were performed as described above with the exception that 32 PCs were used as input for clustering and UMAP. Clustering was performed at different resolutions (0.3, 0.4, 0.6 and 0.8). Clustering at resolution 0.4 resulted in 9 clusters and was in good agreement with the expression of known marker genes for cell types found in brain organoids (Supplementary Fig. 5b). This resolution was therefore selected for delineating different cell populations. Marker genes were identified for each cluster (Supplementary Table 1) using the Wilcoxon rank sum test implemented the FindMarkers function, with a logFC threshold of 0.25 and Bonferroni correction for multiple testing, confirming and refining our initial cell type annotation using canonical marker genes.

### Interfacing cerebral organoid sections with high-density CMOS microelectrode arrays

Whole cerebral organoids were allowed to mature to 4-6 months (see Cerebral organoid generation section) and were then embedded in 10% (w/v) low melting point agarose at ≈ 40 °C and allowed to cool at 2 °C for ≈ 10 minutes. Organoids embedded in the agarose gel were then sectioned into 500 μm thick slices with a vibratome (Leica VT1000S) using an advance speed of one and a vibration speed of nine. The gel block containing the organoid was mounted onto the surface of the vibratome cutting block with super glue and placed into the cutting reservoir filled with ice-cold cutting solution composed of 50% (v/v) DMEM (ThermoFisher), 50% (v/v) Neurobasal (ThermoFisher), 1x penicillin streptomycin (ThermoFisher). The space surrounding the vibratome liquid reservoir was filled with ice to keep the cutting solution temperature constant during sectioning. The vibratome blade was allowed to advance through the front-face of the gel block and organoid tissue, while leaving ≈ 3 mm gel uncut and attached at the back of the gel block. This method allowed previous sections to remain attached to the gel block, which minimized torque on the organoid during subsequent sections and prevented deforming or releasing of the remaining uncut tissue embedded in the gel below. Next, the contiguous layer stack of 500 μm thick organoid sections remaining in the gel block (Supplementary Fig. 1a) was then peeled off using sterile forceps (in a sterile dissection hood equipped with a stereoscopic microscope) and placed into individual wells in a 6 well plate filled with cutting solution using a cut p1000 pipette tip. The slices were then transferred to new set of wells (6 well plate) filled with cutting solution before transferring to individual wells in a 12 well plate filled with BrainPhys neuronal medium (StemCell Technologies) with the addition of the following supplements 2% (v/v) NeuroCultTM SM1 Neuronal Supplement (StemCell Technologies), 1% (v/v) N2 Supplement-A (StemCell Technologies), 20 ng/ml Recombinant Human Brain-Derived Neurotrophic Factor (StemCell Technologies), 20 ng/ml Recombinant Human Glial-Derived Neurotrophic Factor (StemCell Technologies), 1 mM Dibutyryl-cAMP (PeproTech), 200 nM ascorbic acid (StemCell Technologies), 1x penicillin streptomycin (ThermoFisher) and allowed to recover for 24 hours in a 5% CO_2_ incubator at 37 °C prior to placement and positioning on the CMOS MEAs.

High density CMOS MEAs and custom machined (UCSB physics machine shop) liquid reservoir lids made from Delrin (McMaster-Carr), fitted with CO_2_ permeable, water vapor impermeable membranes ^70^, were first sterilized in a 70% (v/v) ethanol solution in deionized water (18.2 MΩ-cm) for 30 minutes. The liquid reservoir of the CMOS MEA was then rinsed thoroughly with sterile ultra-pure distilled water in a sterile hood (≈ 5 ml per CMOS MEA well), and the lids were allowed to air dry in the sterile hood. The recording surfaces of the MEAs were then coated by adding 0.5 ml of poly-l-lysine (PLL) (Sigma Aldrich) solution (0.1 mg/ml in ultra-pure water) into the CMOS MEA reservoirs and then placed in an incubator for ≈ 1 hr. at 37 °C with the lids on. The arrays were then transferred to a sterile hood, the PLL solution was aspirated off and washed an additional 3x with ultra-pure water (1 ml volume for each wash). The CMOS MEA reservoir was then filled with 0.5 ml of BrainPhys medium and the lid was placed on it. The CMOS MEA was transferred to a sterile dissection hood equipped with a stereoscopic microscope. Organoid sections were transferred to the CMOS MEA well with a cut P1000 pipette tip and gently positioned over the recording electrode surface of the CMOS MEA with sterile forceps while visualizing with the stereoscopic microscope (Supplementary Fig. 1b). A sterile custom harp slice grid was used to seat the organoid slice to the CMOS MEA surface. The harp slice grid was made by attaching 122 μm nylon fibers (taken from plastic mesh 9318T45, McMaster-Carr) spaced at a 1 mm pitch to a stainless-steel washer (M3 lock washer, McMaster-Carr) with epoxy (Devcon). The organoid sections on top of the CMOS MEAs were maintained in an incubator (5% CO_2_ at 37 °C), and media was exchanged twice a week.

### Electrophysiology recordings

High density extracellular field potential recordings were measured in a cell culture incubator (5% CO_2_ at 37 °C) using complementary metal-oxide-semiconductor (CMOS) micro-electrode array technology (MaxOne, Maxwell Biosystems, Zurich, Switzerland) containing 26,400 recording electrodes with a diameter of 7.5 μm at a center-to-center distance of 17.5 μm within a sensing area of 3.85 mm x 2.1 mm. A subset of which 1,024 electrodes could be selected for simultaneous recording ^19^. The low-noise amplifiers have a high-pass filter (0.5 Hz) to minimize drift. Electrical measurements were performed once a week (24 hours after a media change), and took around two weeks after sectioning to start spiking with a marked increase in activity in the form of synchronized bursts at ≈ 6 months in age (Supplementary Table 2), reaching a peak in activity at around 7 months (Supplementary Fig. 2a). Automatic activity scans (tiled blocks of 1,020 electrodes) were performed to identify the spatial distribution of electrical activity across the surface of the organoid. Data were sampled at 20 kHz for all recordings and saved as HDF5 file format. We chose the routed top most spiking 1,020 electrodes to have a minimum spacing distance of at least two electrodes (2 x 17.5 μm), providing sufficient electrode redundancy per neuron to enabling accurate identification of singe-units by spike sorting ^21^, while simultaneously resolving network activity across the organoid (Fig. 1).

### Spike sorting

The raw extracellular recordings were automatically spike sorted to extract single-unit activity. The same spike sorting procedure was applied to the planar HD-MEA and the Neuropixels recordings. The raw traces were band-pass filtered between 300-6000 Hz before applying the Kilosort2 algorithm ^20^. The spike sorting output was automatically curated by removing units with an ISI violation threshold ^71^ above 0.3, an average firing rate below 0.05 Hz, and a signal to noise ratio (SNR) below five. For processing data from the same chip with the same electrode configuration, recorded within a short period of time (see stability section in Methods), we concatenated the filtered traces in time before applying spike sorting. All of the processing was performed using the SpikeInterface framework ^21^.

### MEA local field potential signal processing

Raw signals (sampled at 20 kHz) were imported into MATLAB (Mathworks). The local field potential (LFP) component was extracted by first low pass filtering the raw data (frequency cutoff of 500 Hz using 4^th^ order Butterworth filter), and downsampled to 1 kHz. The LFP was then filtered into the traditional oscillatory sub-bands using a bi-directional finite impulse response (FIR) filter ^72,73^. Delta (0.5-4 Hz), theta (4-8 Hz), alpha (8-13 Hz), beta (13-30 Hz) and gamma (30-50 Hz) bands were extracted using *eefilt.m* function from the UCSD EEGlab MATLAB toolbox (Supplementary Fig. 14).

### Acute recordings from cerebral organoids using a high-density Neuropixels CMOS shank

6- month-old cerebral organoids were transferred to BrainPhys media and were cultured for 30 days without shaking before recording. Organoids were then positioned using a cut p1000 pipette tip into a custom well (filled with BrainPhys media) to immobilize the organoid (kept at 37 °C on a temperature-controlled stage). A high-density CMOS shank with 960 electrode sites that tile along a 70 x 20 mm cross-section shank (10 mm long) where 384 electrodes can be programmably routed and recorded from simultaneously ^22^ was inserted into the organoid using a low drift (< 10 nm/hr), motorized precision micromanipulator (MP-285, Sutter Instruments). The Neuropixels probe was mechanically attached to the micromanipulator using a custom machined bracket (UCSB physics machine shop). Electrophysiology data was acquired using SpikeGLX (Bill Karsh, https://github.com/billkarsh/SpikeGLX). Action potential channel was sampled at 30 kHz. The local field (LF) channel (sampled at 2,500 Hz) was low pass filtered at 500 Hz. Downstream processing and analysis were performed with methods similar to 2D CMOS data.

### Single-unit firing rate distributions

The firing rate distributions were calculated directly from the inter-spike interval for each sorted unit as follows: *R_j_*(i) = 1 / Δ*t_i_*, where Δ*t_i_* = (*t_i+1_ – t_i_*) were the time intervals between *i*^th^ subsequent spike events for a given neuron represented by unit *j*. The cumulative distributions of firing rates for a given unit were calculated using the empirical distribution function *ecdf*. The top 80^th^ and 20^th^ percentiles were calculated using the function *prctl* to visualize the bounds of the cumulative distributions across all units. Next, the firing rate cumulative distributions were fit to the non-linear Weibull cumulative distribution function of the form *g_λ,k_* (*R*) = 1 – *e*^(-(*R/λ*)*k*)^, where *R* is the firing rate and *λ* and *k* are the shape and scale parameters, respectively. The function *wlbfit* was used to determine initial conditions for λ and *k* and fed into the nonlinear regression model using *fitnlm*. Iterations from the nonlinear regression model converged to solutions determined by *wlbfit*. R-squared values were determined from the nonlinear regression model. A two-sample Kolmogorov-Smirnov test was performed to assess differences between the nonlinear fit parameters *l* and *k* distributions for a given organoid for control conditions relative to diazepam treatment.

### Population-averaged spike rate and population bursts

For each recording, a population averaged firing rate vector was computed for the spike sorted single-units by summing the spiking activity over all units for each recording frame and averaging these summed values with a 5 ms moving average. For each burst, a burst population rate (5 ms moving average and 5 ms Gaussian) was selected by taking a window with respect to the peak of the multi-unit activity (MUA) (defined below). For each pair of bursts, the similarity in the population rates were assessed by taking the average of the absolute difference between the two burst population rate vectors. This difference was then averaged over all burst pairs. Burst-to-burst similarity between control and diazepam was compared by computing the ratio in the average population rate vector difference between control and diazepam (*ρ_C_* – *ρ_D_*)/(*ρ_c +_ ρ_D_*), where *ρ_C_* and *ρ_C_* are the population rate differences for control and diazepam, respectively. This analysis was performed using a range of time windows with respect to the peak of the MUA activity. Window start times were selected between −200ms until 0ms with respect to the MUA peak with a step size of 10ms. Window end times were selected between 0ms until 500ms with respect to the MUA peak with a step size of 10 ms. The average ratio over all the different time windows was computed to compare the control and diazepam treatments for the different organoids.

Multi-unit activity (MUA) was extracted by band-pass filtering (cutoff frequencies of 0.3 and 7 kHz using a 4^th^ order Butterworth filter) the raw voltage signal, and spike detection was set at 5x the rms-noise for a given electrode ^74^. Detected spike times (rounded to the nearest ms) were then stored as a binary spike time matrix (columns represent electrodes and rows represent time). MUA population-averaged spike rates were calculated by first summing all detected spikes (across all 1,020 electrodes) occurring within a one ms time window to create a *population spike* vector. The *population spike* vector was then averaged over a 20 ms sliding window and smoothed with a 100 ms Gaussian kernel, yielding a population-averaged spike rate as measured across the CMOS array. Next, population bursts peaks were defined when the population-averaged spike rate exceeded 2x its rms value (using the *findpeaks* function with a minimum peak distance of 1 s). Population burst durations were defined by time windows surrounding the burst peak where the burst amplitude was attenuated by 90% of its maximum value.

### Spike time tiling coefficient to draw a functional connectivity map

Correlated spiking activity between single-unit spike trains was compared using the spike-time tiling coefficient (STTC) ^38^. A publicly available MATLAB script ^11^ was used to compute the STTC. A correlation window (Δ*t*) of 20 ms was used to compute a score of functional connectivity weights across all units. The remaining correlations between single unit pairs were further filtered based on the spike time latency distributions of the spike trains. All electrode pairs with a multimodal latency distribution (Hartigans’ dip test *P* < 0.1), a full width at half maximum (FWHM) larger than 15 ms were removed. To rule out the influence of outliers, these measures were computed over all spike time latencies within a [-20;20] ms range. Using the STTC scores of the remaining unit pairs, functional connectivity networks were generated.

### Lower bound connectivity score estimation

To obtain a lower bound for connectivity scores to use for generating connectivity networks, STTC connectivity scores for randomized spike time vectors were computed. Randomization was performed by selecting the spike times within an individual burst event and shuffling the corresponding electrode numbers. This processing was done for each burst event individually and resulted in randomized spike time vectors while maintaining the same population rate as the original recordings. Connectivity scores between electrode pairs were computed in the same way as the original data. Based on the distributions of the STTC scores of the remaining connections, a lower bound threshold of STTC = 0.35 was chosen. Within the randomized data sets (*n* = 6 organoids), 93.2% ± 14.7% percent of the STTC connectivity edges strengths were below the 0.35 threshold across 13 separate measurements; organoid L6 had no pairwise spike correlations (edges) after spike train randomization and was excluded from the statistics above (Supplementary Fig. 10b for randomization distributions).

### Global connectivity network metrics

Global and local connectivity metrics for the connectivity networks were analyzed using the MIT strategic engineering network analysis toolbox for MATLAB ^75^. For networks generated using an STTC, all edges were removed that fell below a threshold of 0.35 (see Lower bound connectivity score estimation detailed above). The size of the largest and second largest component, defined as the number of nodes present in the largest and second largest structure of connected nodes. The size of the second largest component was less than 2% of the size of the largest component for L1, L2, L3 and L4 and less than 10% for L5 and L6 for 7-month-old organoids.

### Directional connectivity map

Based on the mean of the spike time latency distributions, directionality was assigned to the edges in the functional connectivity map. A negative mean spike time latency compared to the reference unit indicated a directionality towards the reference unit and vice versa. Based on the directionality, the in-degree (*D*_in_) and out-degree (*D*_out_) were computed per unit, defined as the number of incoming and outgoing edges respectively. ‘*Receiver’* nodes were defined as (*D*_in_−*D*_out_)/(*D*_in_+*D*_out_) > 0.8, while ‘*sender*’ nodes were defined as (*D*_out_ *-D*_in_)/(*D*_in_+*D*_out_) > 0.8. All other nodes were labelled *brokers*. The directionality per node was calculated for the largest component of interconnected nodes. To evaluate the significance of *sender* and *receiver* nodes, networks were randomized by performing double edge swaps (10 times as many swaps were performed as edges in the control network) and contained the same number of nodes and the same edge density. Randomized networks contained no *sender* or *receiver* nodes.

### Diazepam treatment and analysis

Diazepam (Sigma) was solubilized in DMSO (Sigma) at 1,000 times the target concentration and 1 μL was added to the 1 ml CMOS MEA reservoir containing culture media and gently mixed with a p100 pipette. The cultures (*n* = 4) were then allowed to sit undisturbed for 15 minutes in the incubator (37 °C 5% CO_2_) before recording. Organoid dose responses were carried out over 3, 10, 30 and 50 μM diazepam concentrations (Supplementary Fig. 8a-d).

For connectivity networks generated using recordings from organoids, treated with 50 μM diazepam, the shape of the edge strength distributions in the largest component then were compared. A power law fit (y = a · *x*^b^), an exponential fit (*y* = a · e^-c * *x*^ + d) a truncated power law fit (*y* = a · *x*^b^ · e^-c·^ *^x^*) and a Gamma distribution fit (y = (a/(Γ(b) · c^b^)) · *x*^b-1^ · e^-*x*/c^ were performed on the binned edge strength (pairwise spike train correlations determined by the spike time tile coefficient) distribution (bin size = 0.05). The goodness of the fit was assessed using the Akaike information criterion (AIC) ^46,76^ (Supplementary Fig. 12e).

### Edge change

A gamma distribution was found to be the optimal fit (lowest AIC score, Supplementary Fig. 12e) and used to further assess differences in the distribution of pairwise spike correlation strength determined by the STTC. The fractional change correlation strengths between pairwise spike trains was defined as: (*f*_d_- *f*_c_)/(*f*_d_ + *f*_c_), where *f*_c_ and *f*_d_ are the fraction of total edge strength distribution fits to the gamma distribution for control and diazepam (50 μM), respectively.

Pairwise spike correlation distributions were split into three unique sets of edges. The first set contained edges that were present in both the control and the diazepam recording. The second set contained edges that were silenced by diazepam, meaning that they were only present under control conditions. The third set contained edges that were induced by diazepam, meaning that they were only present under the diazepam condition. Pairwise spike train correlation distributions and connectivity maps were generated for the three different sets of edges.

### Time course in culture analysis

Electrical recordings were made from organoids at 6, 7 and 8 months were compared from three organoids. First, the MUA population-averaged firing rate (defined above) was calculated for each burst period. The results were averaged over all bursts within a recording centered about the population burst peak. Secondly, the peak MUA population rate was averaged over all bursts within a recording and compared between organoids at different ages. The average inter-burst interval, defined as the average time between burst peaks was compared. Functional connectivity maps based on the STTC were generated for the organoids at the different time points.

### Stability analysis

Electrical recordings were made from organoids spaced four hours apart. The number of spike sorted singe-units detected in each recording were compared for two separate organoids (Supplementary Fig. 12a). In addition, pairwise spike train correlation scores were computed between all single-unit pairs. The pairwise spike train correlation distributions for the two measurement time points were compared by a two-sample Kolmogorov-Smirnov test (Supplementary Fig. 12b).

### Theta coherence

Normalized cross-correlation analysis was performed between all pairwise electrodes (1,020) for the theta band time series using the *xcorr* function in MATLAB. The correlation coefficient and respective lag-time were stored in 1,020 x 1,020 matrices. Before performing pairwise correlations, theta oscillation amplitudes smaller than 10 μV were excluded and considered below the noise floor of the CMOS detectors at this bandwidth (Supplementary Fig. 15 for noise floor measurements with synaptic blockers and TTX). Next, each electrode correlation strength was determined by ranking by the summed correlation coefficient between all pairwise electrode sites. The top correlated electrode (‘*seed*’ electrode) was then used to generate a spatial correlation and phase-lag map between the other 1,019 electrodes across the array. This analysis was also repeated for theta time series occurring during population burst periods (see population-averaged firing rate and population bursts in Methods for details). Next, the total number of pairwise theta correlations occurring within the burst periods and during the whole time series were compared. Finally, the total number of pairwise correlations with a correlation score of at least 0.2 was compared between control and diazepam conditions during burst time periods.

To further validate the cross-correlation calculations, theta oscillations were signal averaged relative to the ‘*seed*’ electrode. First, positive peaks were identified using the *findpeaks* function with a minimum peak height of 10 μV (see Supplementary Fig. 15e and Blockers and TTX analysis in Methods) and a minimum peak distance of 100 ms. Next, the theta peak time points from the ‘*seed*’ electrode were used as anchor points to average theta signals centered over a 500 ms time window for each electrode. The amplitude and relative phase shifts (relative to the ‘*seed*’ electrode) were used to generate spatial coherence maps of theta oscillations across the organoid.

### Burst peak-centered theta signal averages

Event triggered averages were made of the theta filtered LFP from each electrode using the peak of the population burst as the anchor point. For each electrode the theta filtered signal was averaged over all burst events. Spatial maps were generated showing the theta phase per electrode at different time points relative to the population burst peak. The amplitude of the envelope of the averaged theta signal was used as a measure for the consistency of the theta phase relative to the MUA population burst peak for multiple burst events. Similarly, the MUA population burst peak centered theta phase was obtained for each burst of spiking activity (over a 5 ms sliding window). Theta phase angle spread was determined by calculating the circular standard deviation of the theta phase relative to the population burst peaks ^77^. The circular standard deviation (defined as the phase angle spread in Fig. 7g) over all bursts of spiking activity was computed for each time point relative to the burst peak. For each electrode where the circular standard deviation fell below one within 250 ms before the burst peak until 500 ms after the burst peak, the time point relative to the peak with the lowest circular standard deviation was obtained. Using these relative time points of lowest circular standard deviation, a histogram showed the time relative to the burst peak with the most consistency in the theta phase of the signals taken over all individual spiking activity bursts.

### Theta activity within and outside of neuronal population bursts

Cross-correlation analysis of the theta band time series, as described above, was performed on the whole time series and as cross correlations on theta time series occurring within population bursts (detailed in Population-averaged spike rate and population bursts section of Methods). The number of correlation scores of at least 0.2 was compared between whole theta time series data and the theta time series occurring during population bursts. Next, for each electrode that had a correlation score of at least 0.15 with the reference site for the population burst time series, the envelope of the theta filtered LFP was generated and the rms of the envelope was computed. The fraction of the recording for which the envelope exceeded 1.5 times the rms was computed for the recording frames within burst periods and the recording frames outside burst periods. The number of electrodes that showed at least a 10% increase in the fraction of time during which the envelope was above the rms threshold during bursts periods compared to non-burst periods was counted. Similarly, the number of electrodes with a 10% increase during non-burst periods compared to burst periods was counted.

### Imaginary coherence estimation

The raw LFP signal, band-pass filtered into the theta band was used to compute the degree of coherence using the imaginary part of the coherence estimation. The imaginary coherence metric between each of the electrode pairs ^50^ was computed using a window function with the length of 0.5 s and 25% overlap. The connectivity strength at each electrode, regional imaginary coherence, was estimated by averaging across all Fisher’s Z-transformed imaginary coherence values ^78^. After generating the imaginary coherence matrix for the full 1,020 x 1,020 electrode pairs, we thresholded this data-matrix at the 90^th^ percentile of the imaginary coherence values. We selected the nodes with >200 connections in this limited matrix, which resulted in seven highly coherent electrodes (Supplementary Fig. 18b). These seven electrodes were connected to 595 other electrodes in total. These nodes were clustered based on squared Euclidean distance and k-means clustering method (*evalclusters* function in MATLAB). The clustering algorithm with its highest stability, identified two clusters, and separated a small cluster with 54 electrodes, which was locally distributed on the array (cluster 1; Supplementary Fig. 18c) and had tighter within coherence (Supplementary Fig. 18e). In three different organoids, for each of the 1,020 electrodes, the average strength of the imaginary coherence was calculated as the row-averages for the 1,020 x 1,020 matrix, before and after treatment with diazepam. Paired t-tests comparing the pre- and post-diazepam conditions for each of the organoids showed a significant reduction of imaginary coherence with diazepam treatment (Supplementary Fig. 18f).

### Blocker and TTX experiments and analysis

Preparation of stock solutions to block components of fast synaptic transmission: The AMPA receptor blocker NBQX (Abcam) was solubilized in DMSO, the NMDA receptor blocker R-CPP (Abcam) and the GABA receptor blocker Gabazine (Abcam) were prepared in ultrapure distilled water (Life Tech) at 1,000x the desired working concentration. The working concentrations were 10, 20 and 10 μM for NBQX, R-CPP and gabazine respectively. The sodium channel blocker tetrodotoxin (TTX) citrate (Abcam) was solubilized in ultrapure water at 1,000x the working concentration (1 μM). Recordings were made from organoids silenced with synaptic blockers and TTX. Spatial maps of the median firing rate per electrode were generated from organoids treated with synaptic blockers, synaptic blockers in combination with TTX and for the control recordings of untreated organoids. For each treatment, the number of actively spiking electrodes was counted. This was defined as having at least 30 spikes over the 120 s window.

For each electrode on the array, the theta oscillation amplitude envelope was computed^79^. The rms of the theta amplitude envelope was calculated for recordings of the organoids silenced with synaptic blockers and TTX. Subsequently, the fraction of the total recording time that the theta envelope exceeded a threshold of 1, 1.5 or 2 times the rms was computed for each electrode, as well as for the burst periods and the non-burst periods of the same electrode during the control recording. This fraction of time above the threshold during either the burst period or the non-burst period of the control recording was compared to the blockers and TTX recording. The percent change with respect to the blockers and TTX recording was computed and averaged over all electrodes that spiked at least 30 times during the control recording. Similarly, signal envelopes were obtained for the delta, theta, alpha, beta and gamma frequency bands. For each frequency band in each electrode, the rms was computed from the organoids silenced with synaptic blockers and TTX. The fraction of time that the envelope exceeded 2x the rms was computed for the whole time range of the recording treated with blockers and TTX, as well as for the burst periods of the control recording. The percent change of the fraction of time that the envelope exceeded the threshold was computed for each frequency band in each electrode. In addition, the constant decrease in the LFP amplitude over all frequencies after treatment with blockers and TTX was further demonstrated by comparing the power spectral density plots of the same electrode during control recordings and during recordings after treatment with blockers and TTX. The LFP power spectrum scaled inversely with frequency (∼1/*f*) ^80^ and was above the electronic noise floor in control conditions (Fig S15a-d). The mean theta rms voltage fluctuations measured across all electrode for *n* = 4 organoids was 7 ± 2 μV (mean ± STD) for blocker + TTX experiments (Supplementary Fig. 15e), which is in agreement with noise thresholds at this bandwidth for this CMOS architecture in physiological environments ^19^.

### Spike phase locking to theta

The time series theta phase, *ϕ*(*t*), was determined by taking the standard Hilbert transformation, *H*(*t*), of the theta band oscillation and calculating the angle between the real and imaginary components of *H*(*t*) in MATLAB ^79^. The theta phase was then counted at each spike event time for all spike-sorted units. The Rayleigh criterion was used to test the non-uniformity of spikes distributed across the circular phase angles (0°, 360°) ^55,77,81^. Spikes were considered to be phase-locked to theta if they passed the Rayleigh criteria test for non-uniformity (*P* < 0.05).

