## Supplementary Figures for "Human brain organoid networks"

**This PDF file includes:**

Supplementary Figs. 1 to 20

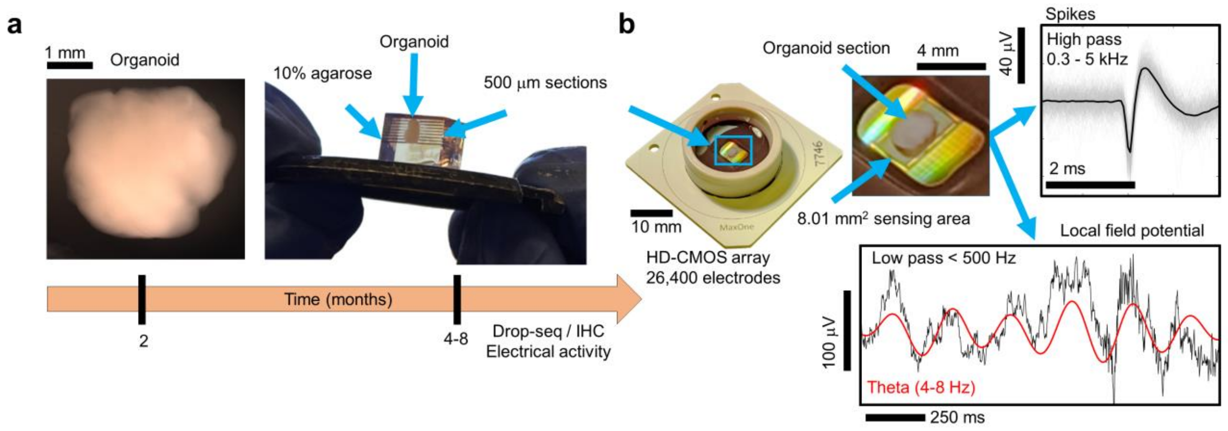


**Supplementary Fig. 1. Human cerebral organoid preparation for electrophysiology. a** Left, picture of a cerebral organoid at 2 months, scale bar is 1 mm. Right, organoids (4-6 months old) were imbedded in 10% agarose and sectioned into 500 µm thick slices. **b** Organoid slices were then placed on high-density CMOS microelectrode arrays (26,400 recording electrodes spanning across surface area of ≈ 8.01 $mm^2$ with a center-to-center distance of 17.5 µm). High bandwidth recordings reveal the presence of high frequency (≈ 1 ms) extra-cellular action potentials (spikes) as well as low-frequency components of the local field potential in the theta range (4-8 Hz), see Methods and Supplementary Fig. 2 for characterization of electrical activity.

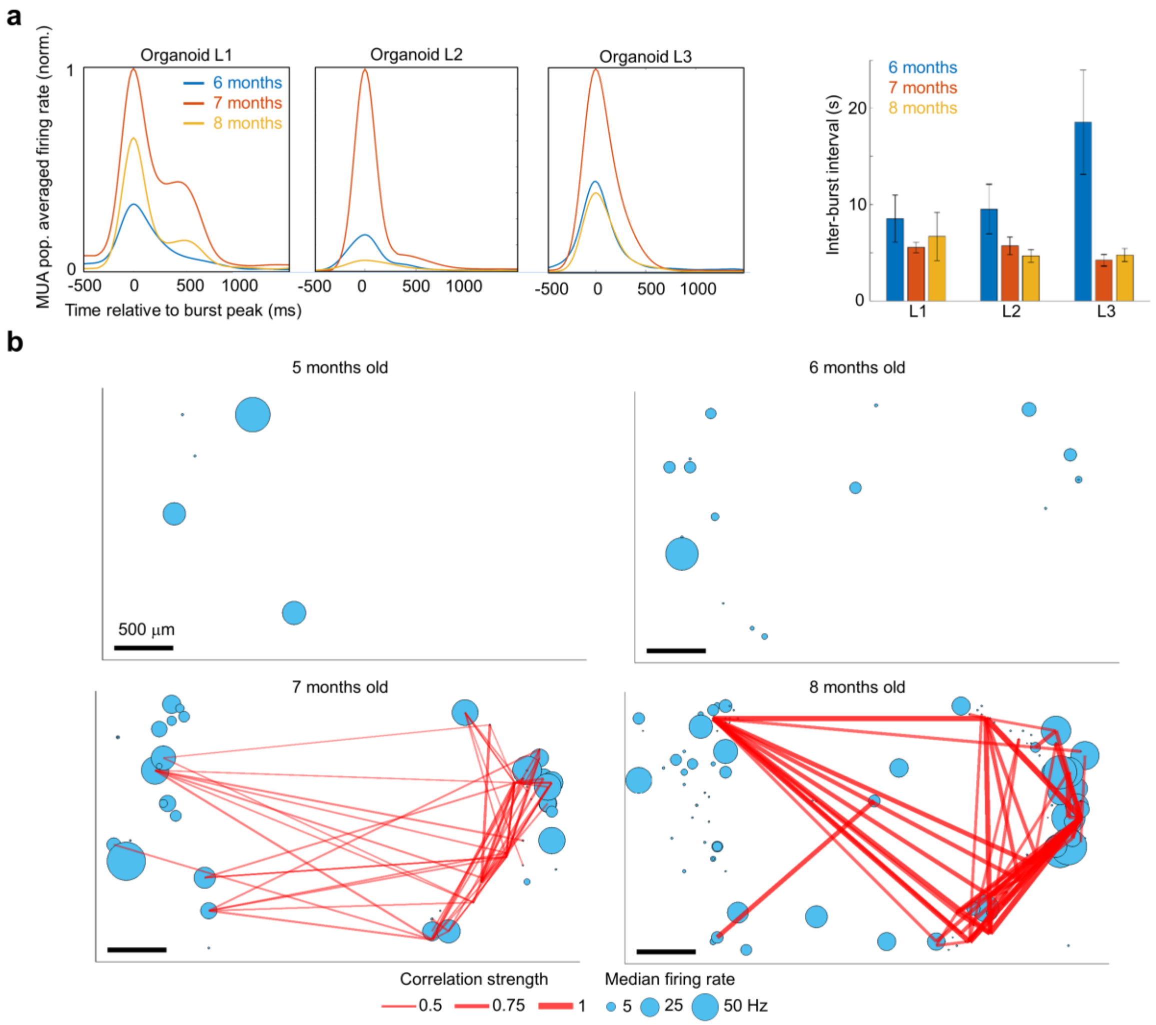


**Supplementary Fig. 2. Overview of organoid spiking activity at different time points on arrays. a** Left, the multi-unit activity (MUA) population-averaged firing rate (pop. rate) is shown at different developmental time points for *n* = 3 organoids and shows a peak in activity at 7 months. Right, the mean inter-burst intervals are shown. **b** Median firing rate from single-unit activity (blue circles) and pairwise spike correlation strengths (red lines) are shown for organoid L1 at different developmental time points (5.1, 6, 7 ,8.1 months old).

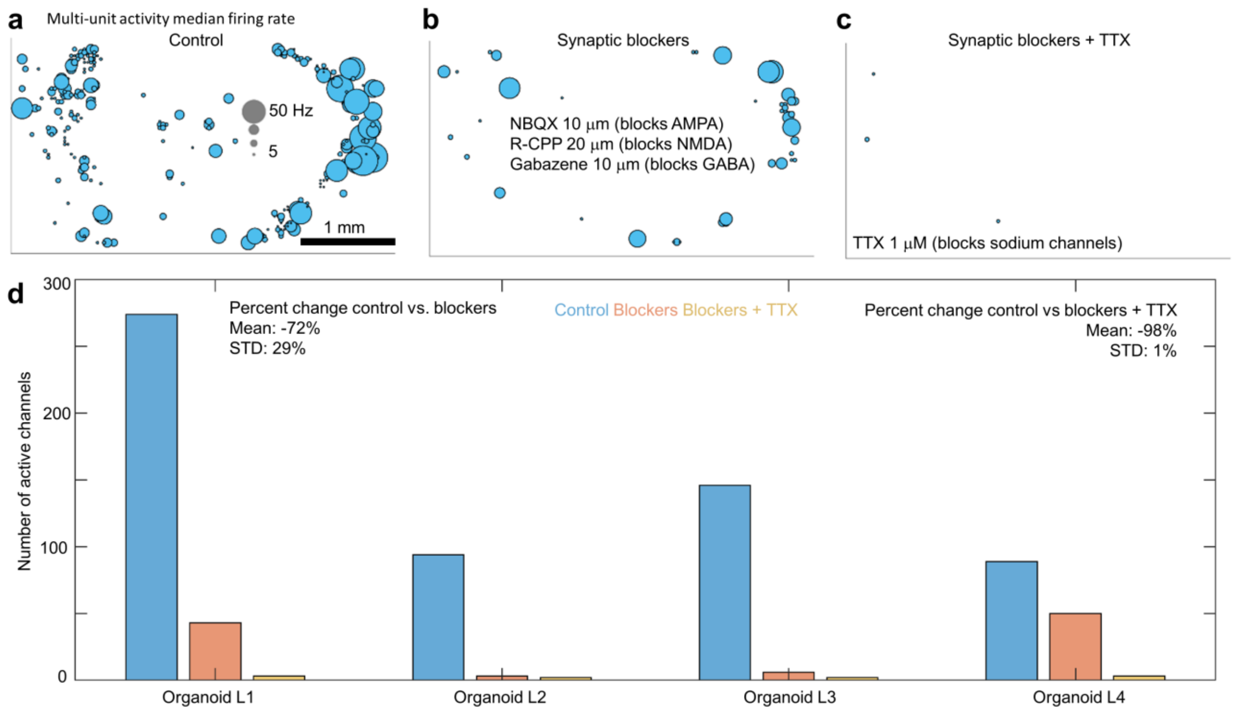


**Supplementary Fig. 3. Synaptic and sodium channel blockers. a-c** The median firing rate calutated from multi-unit spiking activity is shown. The addition of synaptic blockers reduced spiking activity by 72% ± 29% (mean 1 ± STD) at saturating concentrations 10 μM NBQX (blocks AMPA), 20 μM R-CPP (blocks NMDA) and 20 μM Gabazine (blocks $GABA_A$). The addition of 1 μM tetrodotoxin (TTX) reduced detected spikes by 98% ± 1% (mean 1 ± STD) across $n$ = 4 organoids. The remaining small number spike events can be attributed to false detected spikes due to electronic noise. **d** The number of active electrode (with at least 30 spikes) are shown across $n$ = 4 organoids for the same conditions shown in **a-c**.

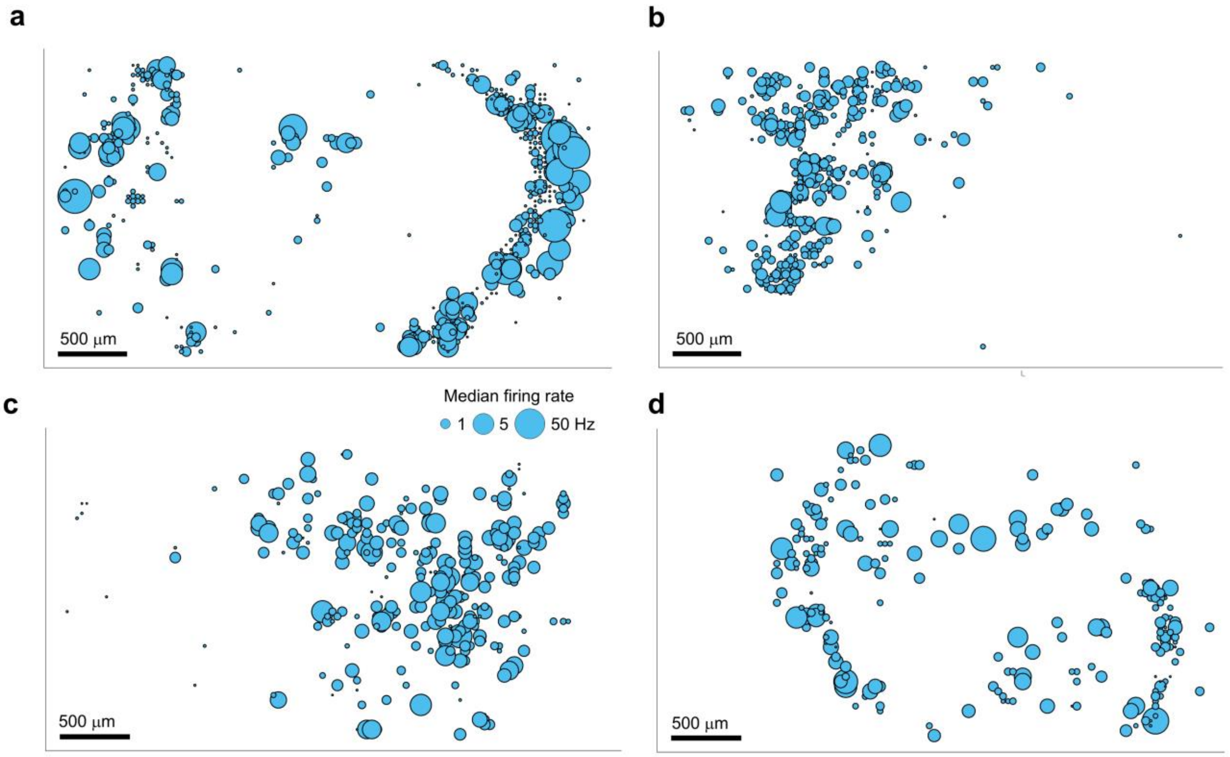


**Supplementary Fig. 4. Overview of spatial spiking patterns across organoids.** The median firing rate measured by individual electrodes (multi-unit activity) are shown across the organoid. **a** Organoid L1 at 7 months old. **b**, **c** Organoid L2 and L3 at 8 months, respectively. **d** Organoid L5 at 6 months old.

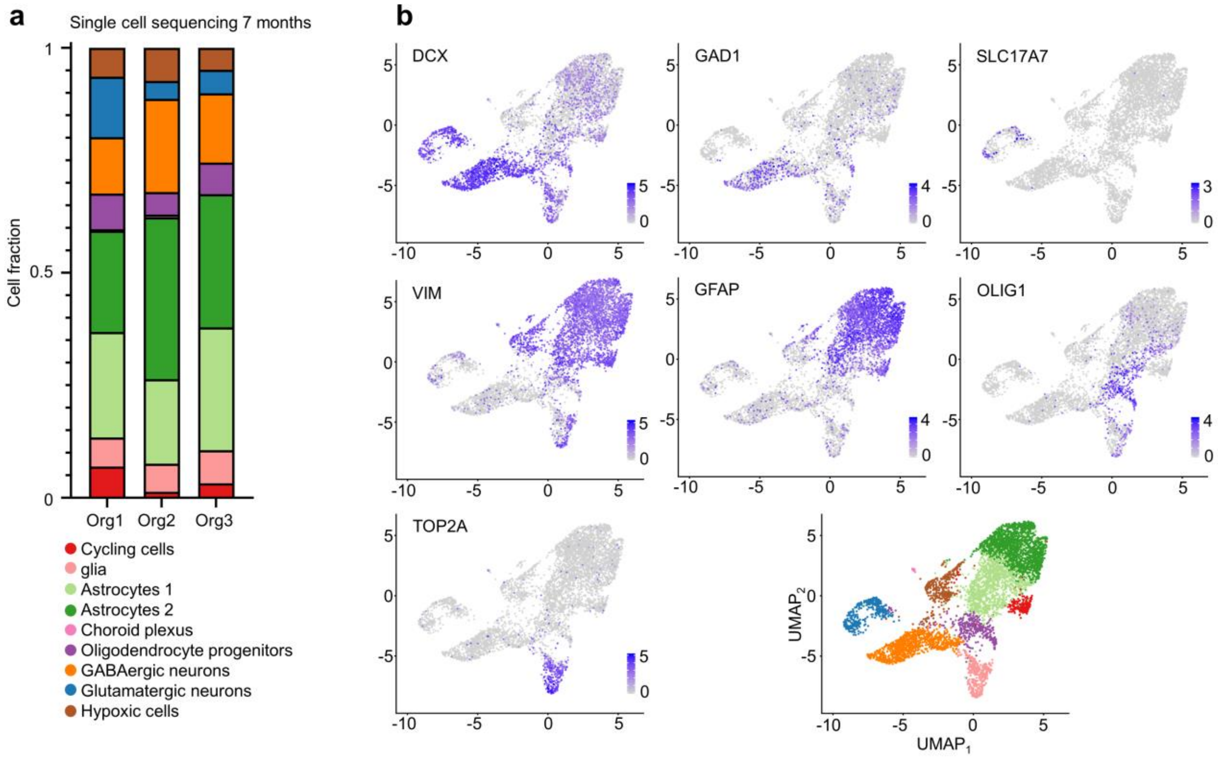


**Supplementary Fig. 5. Human cerebral organoid identification of cell types using single-cell sequencing. a** Relative proportions (fractions) of cell types identified in the 3 individually sequenced 7-months-old organoids as determined by drop-seq. **b** Selection of canonical markers used to identify cell types in single-cell data. UMAP plots are colored by normalized and scaled expression values. DCX: neurons; GAD1: GABAergic neurons; SLC17A7: glutamatergic neurons; VIM: glia; GFAP: astrocytes; OLIG1: oligodendrocyte progenitors; TOP2A: cycling cells. Bottom right UMAP plot is colored by cluster.

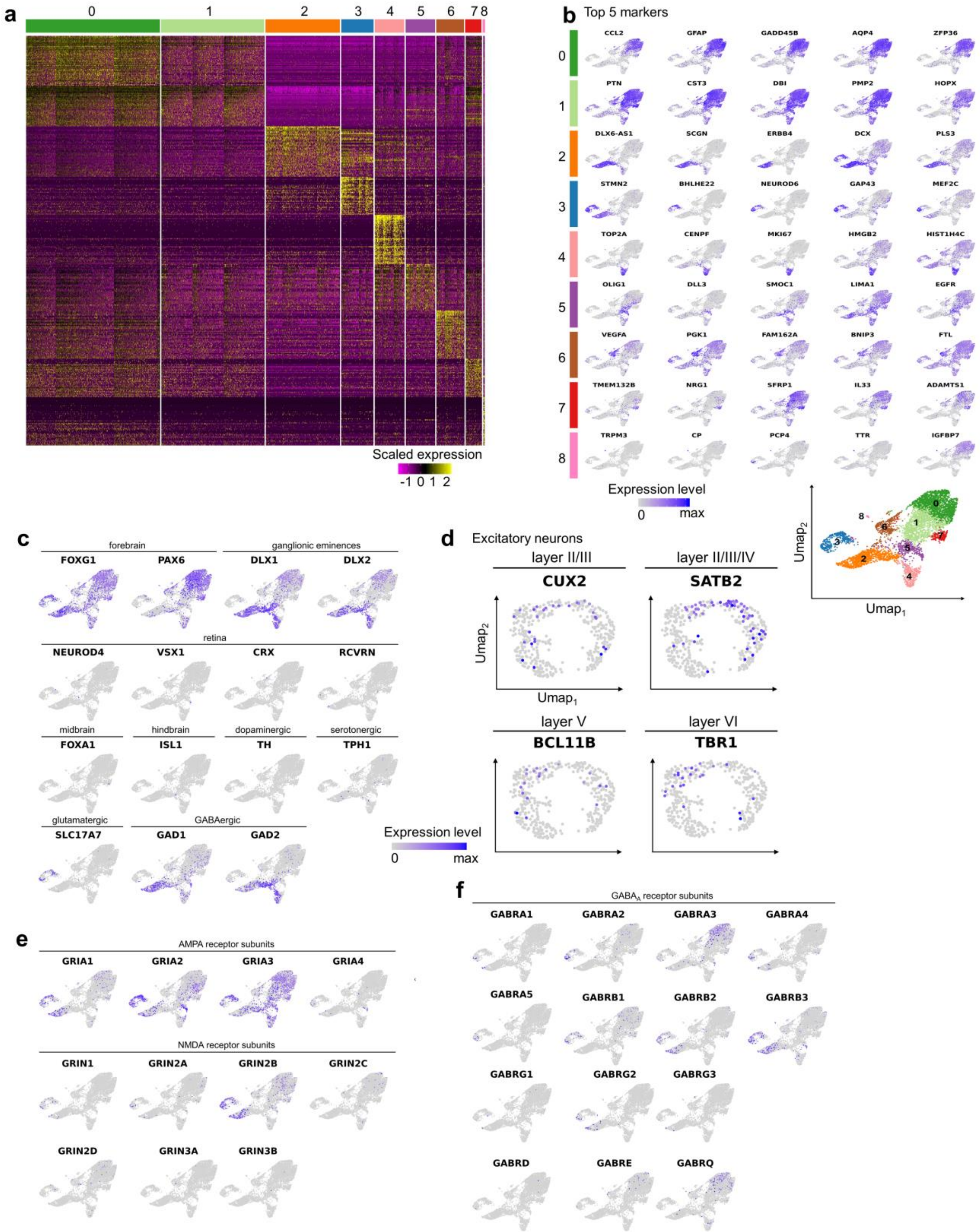
a
0 1 2 3 4 5 6 7 8
Scaled expression
-1 0 1 2
b
Top 5 markers
CCL2 GFAP GADD45B AQP4 ZFP36
PTN CST3 DBI PMP2 HOPX
DLX6-AS1 SCGN ERBB4 DCX PLS3
STMN2 BHLHE22 NEUROD6 GAP43 MEF2C
TOP2A CENPF MKI67 HMGB2 HIST1H4C
OLIG1 DLL3 SMOC1 LIMA1 EGFR
VEGFA PGK1 FAM162A BNIP3 FTL
TMEM132B NRG1 SFRP1 IL33 ADAMTS1
TRPM3 CP PCP4 TTR IGFBP7
Expression level
0 max
$Umap_2$
$Umap_1$
c
forebrain
ganglionic eminences
FOXG1 PAX6 DLX1 DLX2
retina
NEUROD4 VSX1 CRX RCVRN
midbrain hindbrain dopaminergic serotonergic
FOXA1 ISL1 TH TPH1
glutamatergic GABAergic
SLC17A7 GAD1 GAD2
d
Excitatory neurons
layer II/III
CUX2
layer II/III/IV
SATB2
layer V
BCL11B
layer VI
TBR1
Expression level
0 max
e
AMPA receptor subunits
GRIA1 GRIA2 GRIA3 GRIA4
NMDA receptor subunits
GRIN1 GRIN2A GRIN2B GRIN2C
GRIN2D GRIN3A GRIN3B
f
$GABA_A$ receptor subunits
GABRA1 GABRA2 GABRA3 GABRA4
GABRA5 GABRB1 GABRB2 GABRB3
GABRG1 GABRG2 GABRG3
GABRD GABRE GABRQ

**Supplementary Fig. 6. Cellular identities and neurotransmitter profiles of cerebral organoids determined by single-cell transcriptomics. a** Heatmap of the top 50 marker genes for each cluster, colored by z-scaled expression values. **b** Expression patterns of top 5 marker genes for each cluster. UMAP plots show expression levels (normalized and scaled) of the top 5 marker genes for each cluster. **c** Expression patterns of brain-region specific marker genes reveal predominant forebrain identity of the cerebral organoids used in this study. **d** Expression of pyramidal layer markers in glutamatergic neurons. Glutamatergic neurons were extracted from the dataset and projected as UMAP. Plots are colored by expression of selected cortical layer markers. **e** Organoids express multiple genes encoding subunits of AMPA- and NMDA-types of glutamate receptors. UMAP plots are colored by expression level of all known genes encoding subunits of AMPA and NMDA receptors. **f** Organoids express multiple genes encoding $GABA_A$ receptor subunits. UMAP plots are colored by all known and detected genes encoding $GABA_A$ receptor subunits. Not detected in single-cell data were GABRA6 and GABRP.

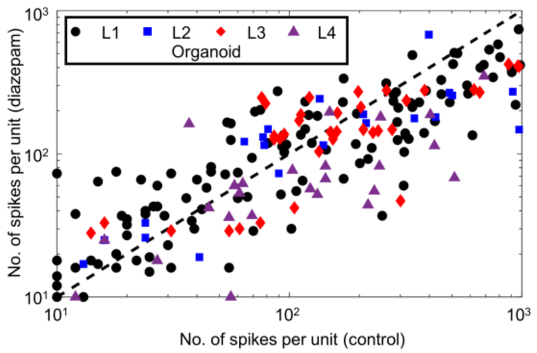


**Supplementary Fig. 7.** Total number of single-unit spikes under control conditions plotted against the number of single-unit spikes under diazepam (50 µM) conditions remain similar across $n$ = 4 organoids.

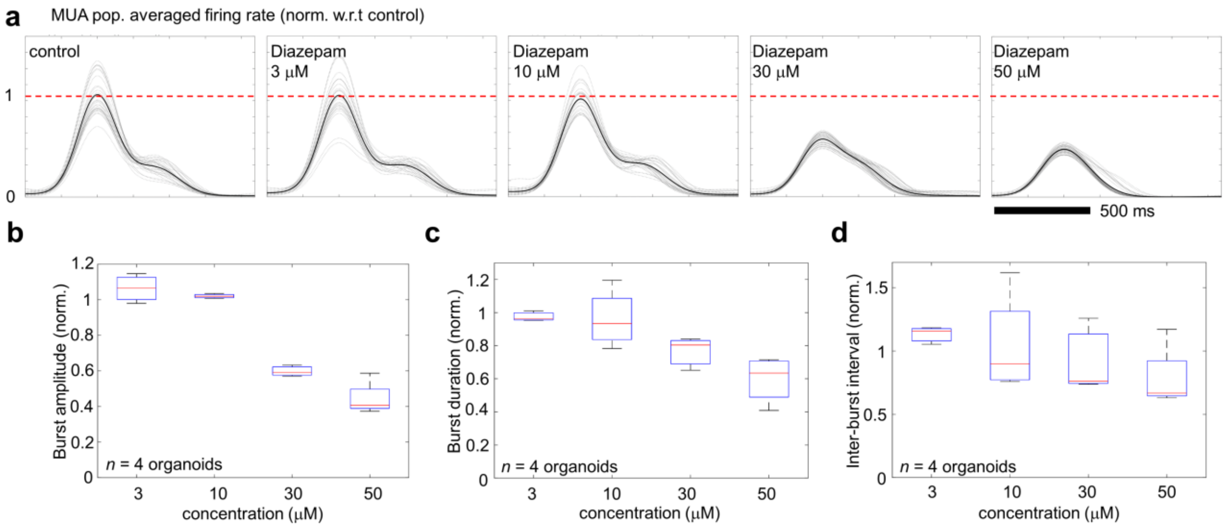


**Supplementary Fig. 8. Pharmacological effects on spiking activity. a** Multi-unit activity (MUA) population averaged firing rate (20 ms average sliding window smoothed with a 100 ms Gaussian kernel) for organoid L1 at 7.7 months old for control (left) and increasing concentrations of diazepam (3, 10, 30, 50 μM) measured over 120 s. The gray lines are MUA population rates plotted for each burst (relative to the burst peak), the black line is the average across all bursts. **b-d** Mean burst amplitudes, burst durations and inter-burst intervals are reduced for diazepam concentrations greater than 10 μM for *n* = 4 organoids. Values are normalized with respect to control conditions.

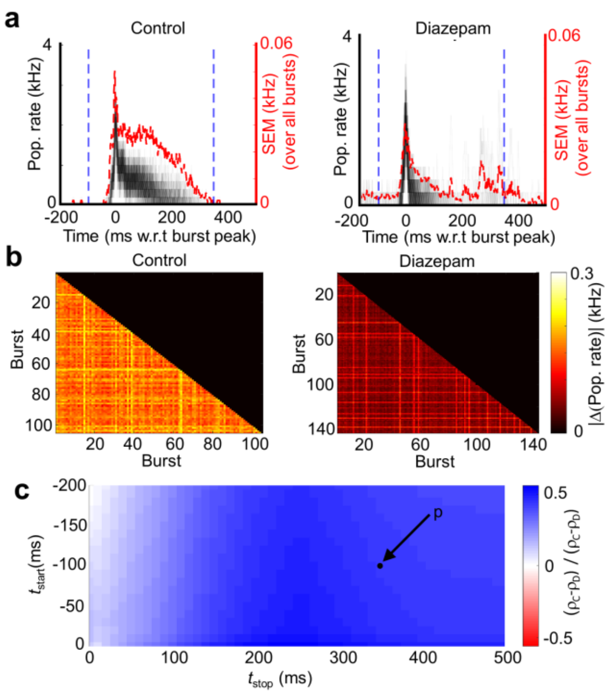


**Supplementary Fig. 9. Variation in population-level dynamics for organoid L2. a** The population averaged firing rate for the control and diazepam recordings calculated from single-unit activity averaged over a 5 ms window. The population rate for each burst is plotted individually, centered by the peak in multi-unit activity (MUA). The standard error of the mean (SEM) calculated over all bursts is plotted in red. **b** The average difference in the population rate is shown across all individual burst pairs for the control and diazepam (50 µM) recordings from the same organoid. The average population rate difference is computed over a time window of -100 ms until +350 ms relative to the MUA peak (blue dotted line in **a**). The color bar scale is the same for control and diazepam. **c** Fractional change between population rate for control and diazepam $(\rho_C - \rho_D)/(\rho_C + \rho_D)$, averaged over a range of different time windows for L2. Here, $\rho_C$ and $\rho_C$ are the average difference in the population rate (for a given time interval $t_{start}$ and $t_{stop}$) for control and diazepam, respectively. Blue tiles indicate a higher average population vector difference for control. The black dot ($P$ = (-100, 350 ms)) indicates the time window used in **b**.

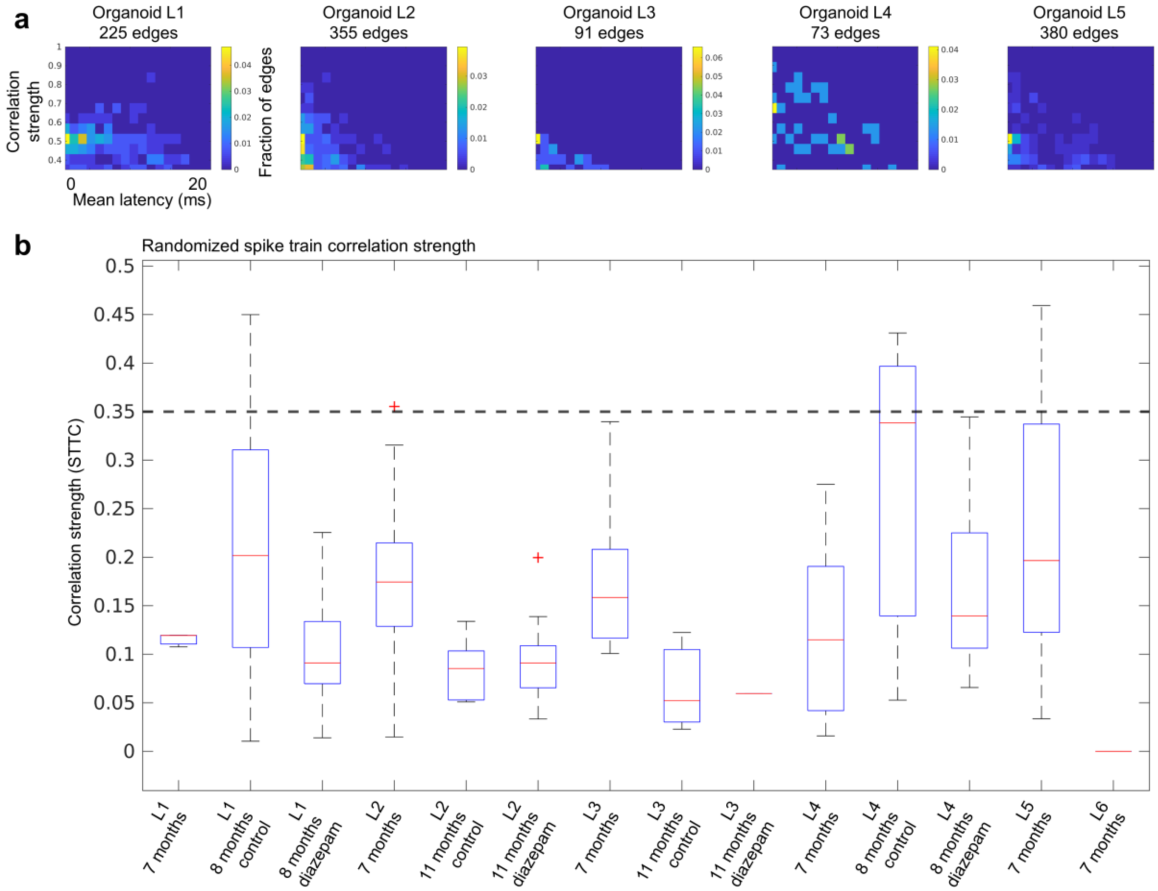


**Supplementary Fig. 10. Latency distributions. a** Pair-wise spike latency distributions as a function of correlation strength for *n* = 5 organoids at 7 months old consisting of unimodal latency distributions with a FWHM < 15 ms. Multimodal latency distributions (Hartigans' dip test *P* < 0.1) were excluded. Organoid L6 is not plotted due to low number of edges. **b** Connectivity edge strength distributions for randomized spike data from six organoids are shown for multiple measurements (see Methods); 93.2% ± 14.7% percent of the connectivity edges strengths were below the 0.35 threshold (*n* = 6; 13 separate measurements). Organoid L3 at 11 months for diazepam treatment only had one edge while L6 had no pairwise spike correlations (edges) after spike train randomization and was excluded from the statistics above.

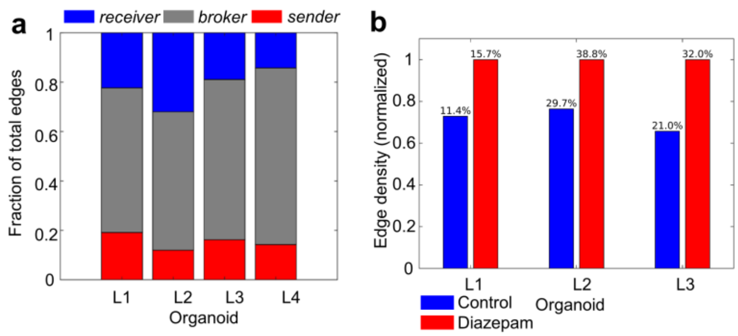


**Supplementary Fig. 11. Diazepam modulation of organoid connectivity. a** Bar plots indicate that the relative fraction of '*sender'*, *brokers*' and '*receivers*' nodes remain similar across multiple organoids ($n$ = 4). **b** The edge density between the largest component of interconnected nodes increases for diazepam (50 µM) relative to control. For visual clarity the plots are normalized relative to control conditions. The edge density percentile of the observed network is shown above the bar plots.

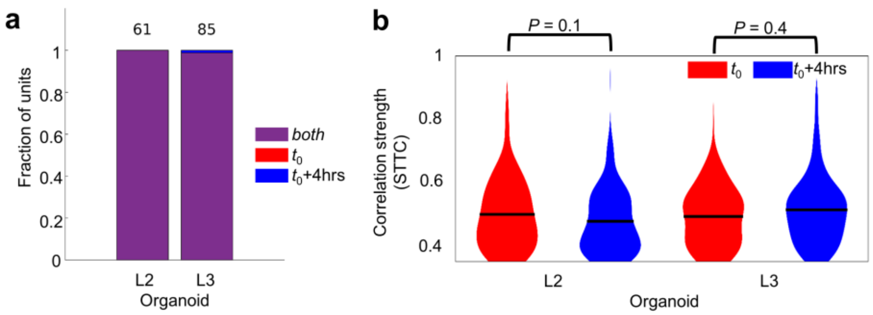


**Supplementary Fig. 12. Stability of active units. a** The number of active spiking units remain similar for two separate recordings taken at time $t_0$ and $t_0$ + 4 hours for two organoids (8-months old). Purple bars indicate the number of units active for both recordings, the red and blue bars represent the number of unique units at time $t_0$ and $t_0$ + 4hrs, respectively. **b** The distribution of edge strengths determined from the spike time tile coefficient are not significantly different as determined by two-sample KS test ($P \geq 0.1$) when comparing the initial recording at time $t_0$ (red) and time $t_0$ + 4 hours (blue) for the same two organoids. The number of edges for time point $t_0$ are 85 and $t_0$ + 4 hrs are 154 for organoid L2 and 120 and 212 for organoid L3, respectively.

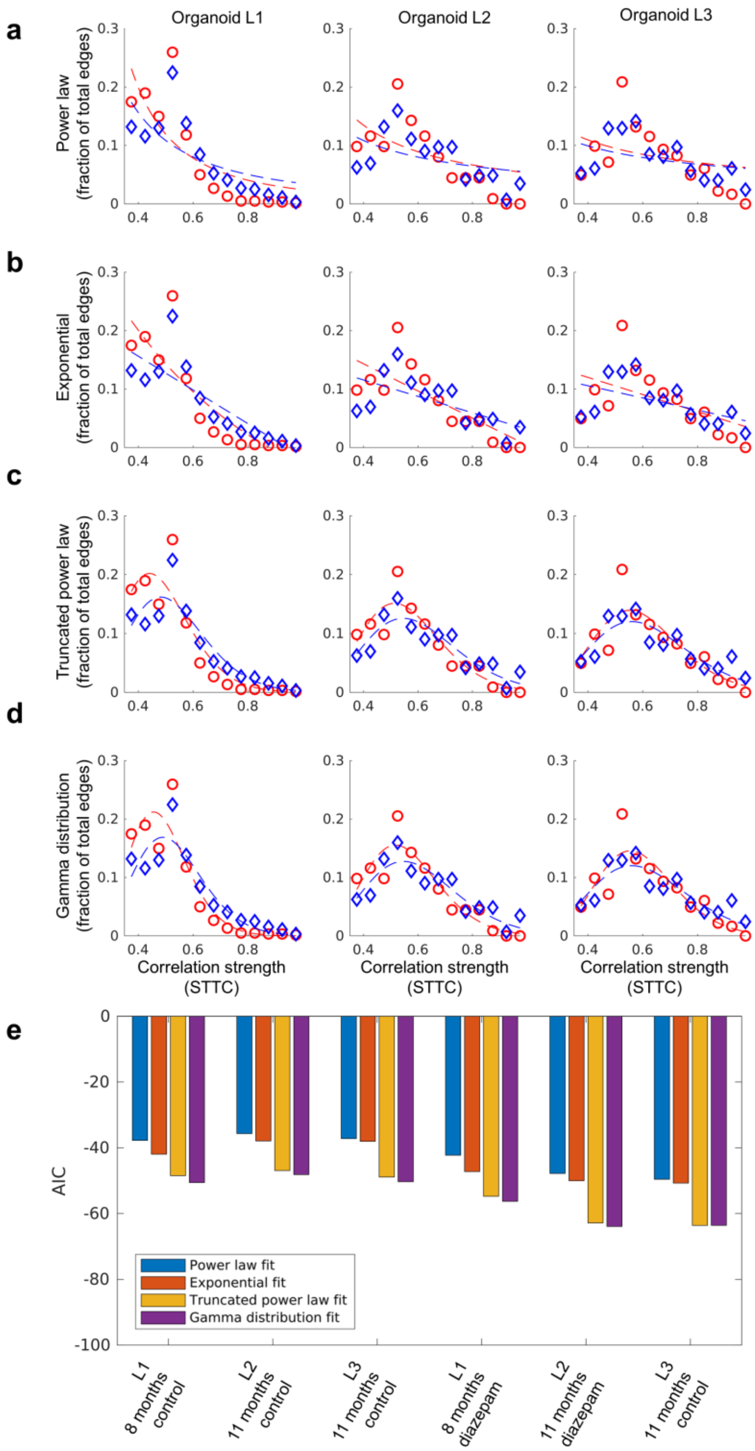


**Supplementary Fig. 13. Edge strength scaling relationships details for control and diazepam data. a** Power law ($y = \mathrm{a}\cdot x^{\mathrm{b}}$). **b** exponential ($y = \mathrm{a}\cdot \mathrm{e}^{-\mathrm{c}x} + \mathrm{d}$). **c** truncated power law ($y = \mathrm{a}\cdot x^{\mathrm{b}}\cdot \mathrm{e}^{-\mathrm{c}x}$). **d** Gamma distribution ($y = (\mathrm{a}/(\Gamma(\mathrm{b})\cdot \mathrm{c}^{\mathrm{b}}))\cdot x^{\mathrm{b}-1}\cdot \mathrm{e}^{-x/\mathrm{c}}$). Fits to the edge strength distributions (edge strengths determined by the spike time tile coefficient (STTC)) were performed on the largest components of the functional connectivity networks. **e** Akaike information criterion (AIC) (see Diazepam comparison section of Methods for details) of the fitted data indicates that the Gamma distribution results are the best fit of the edge strength distributions for $n$ = 3 organoids.

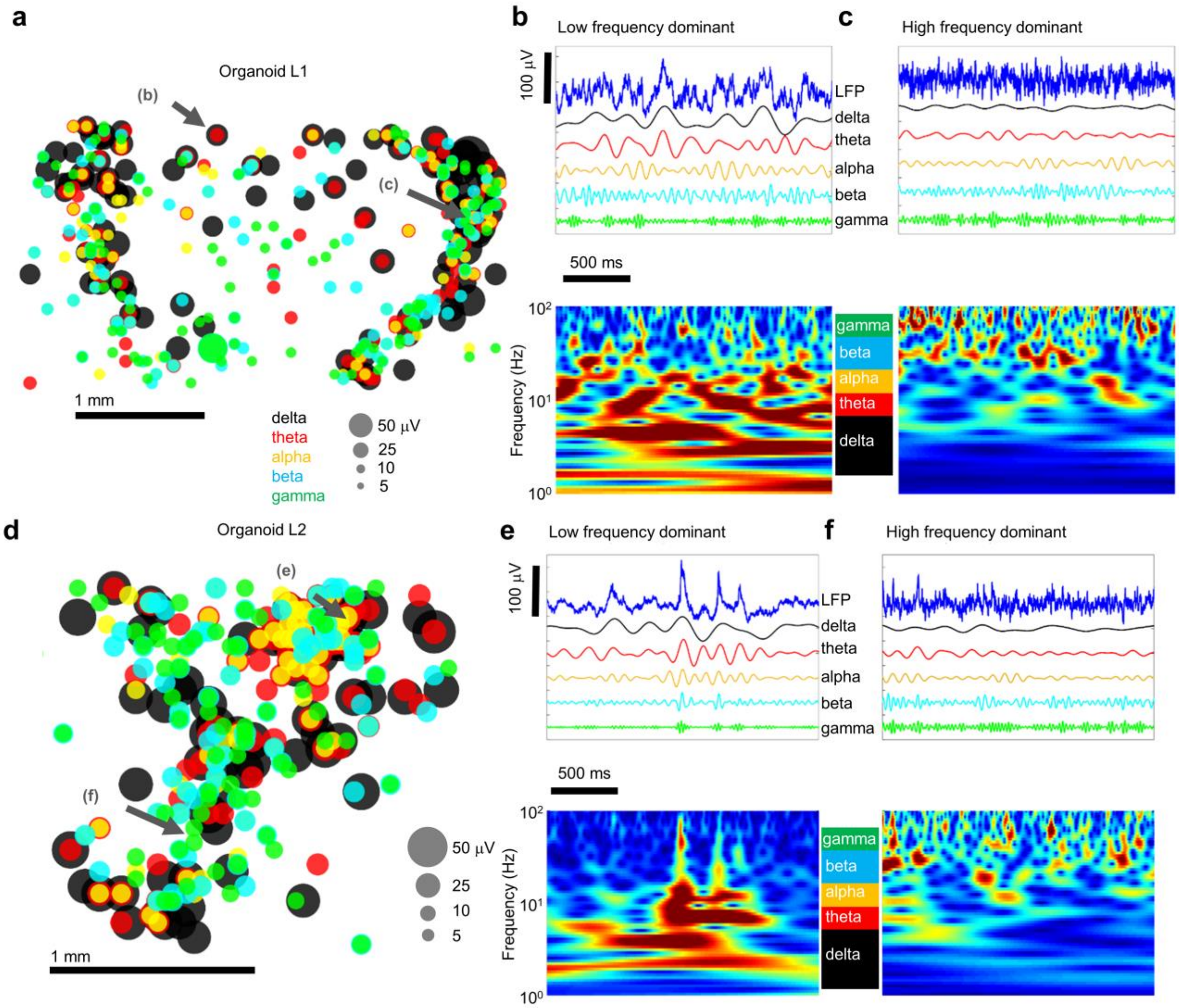


**Supplementary Fig. 14. Overview of LFP bands: spatial amplitudes, sub-band filtering and spectrograms. a** Amplitude map of local field potential (LFP) sub-bands, the top ten percent per band are shown by amplitude: delta (0.5-4 Hz), theta (4-8 Hz), alpha (8-13), beta (13-30) and gamma (30-50 Hz). **b** Top, electrode site with dominant low frequency component of the LFP. Bottom, spectrogram of the LFP is shown. **c** Same as **b** for an electrode with a dominant high frequency component of the LFP. Organoid L1 at 8 months old. **d-f** Same as previous panels for organoid L2 at 8 months old.

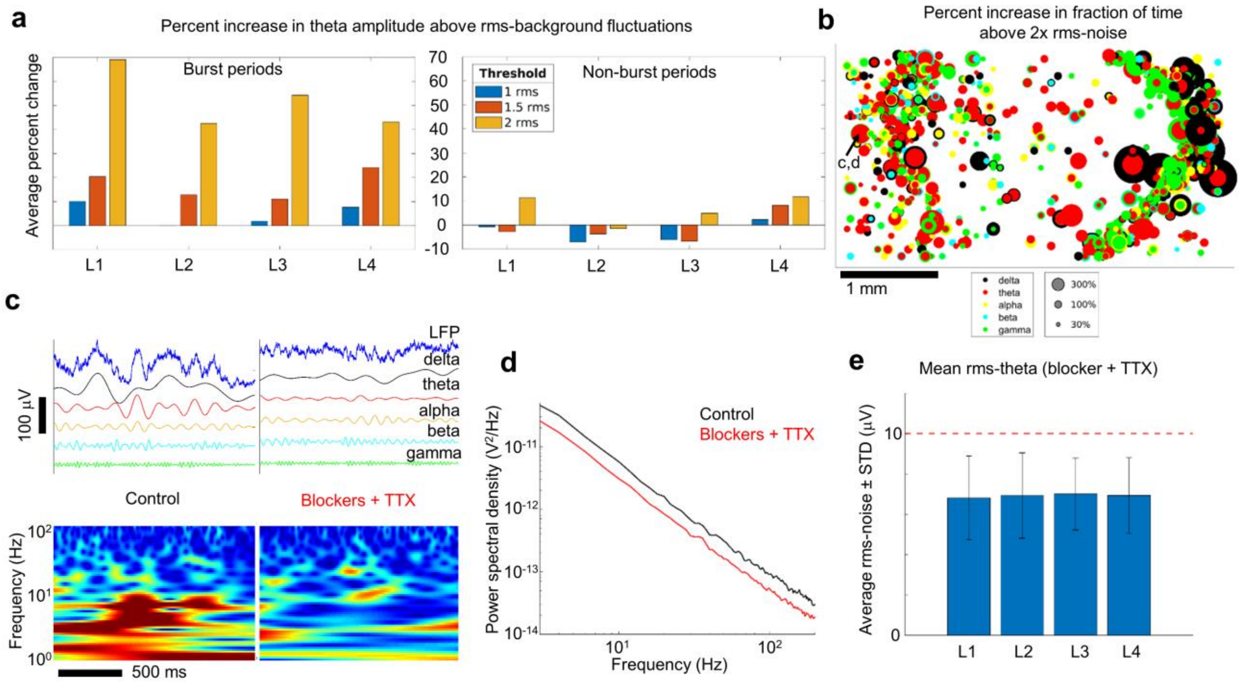


**Supplementary Fig. 15. Synaptic blockers and TTX attenuate LFP amplitude.** **a** Left, theta oscillation amplitude spent more time above the noise floor during population bursts compared to the amplitude measured with synaptic blockers + TTX. Right, there is no observable difference between theta amplitude time spent above the noise floor during non-burst periods compared to blockers + TTX (*n* = 4 organoids). **b** The fraction of time that the signal amplitude falls above 2 times the noise rms for burst periods compared to blockers + TTX recordings shows increased amplitudes in all frequency bands, dispersed over the whole array (organoid L1 at 8 months). **c** Example of the raw and decomposed signal of the channel specified in **b** during the control recording and the recording of the organoid treated with blockers and TTX. **d** The power spectral density is shown for the LFP of the channel specified in **B** shows a decrease in spectral power for all frequencies after treatment with blockers and TTX. **e** The empirical theta LFP noise floor threshold of 10 µV lies roughly 1.5 STD above the rms theta band noise levels measured across *n* = 4 organoids treated with blockers and TTX.

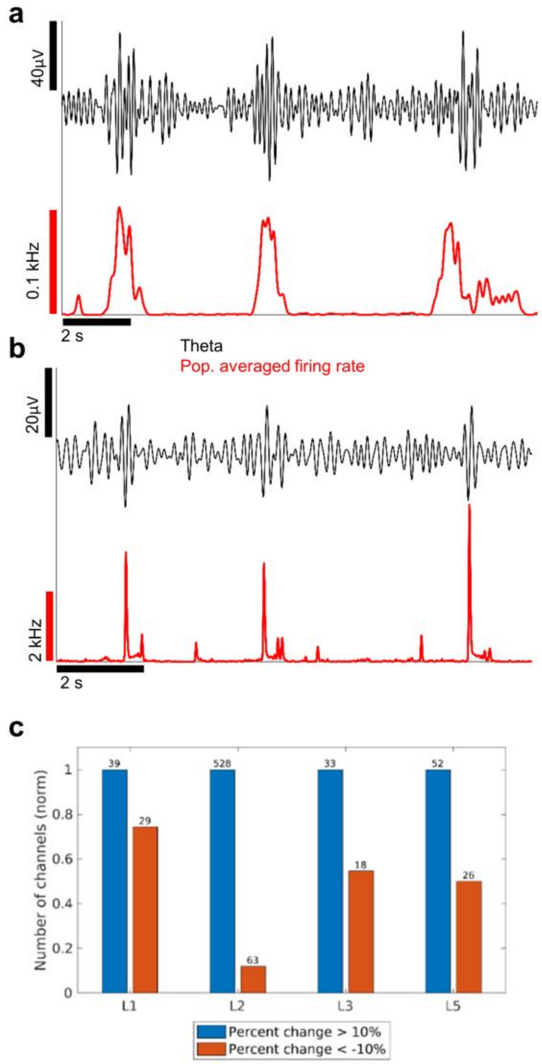


**Supplementary Fig. 16. Theta amplitude within and outside of population bursts. a**,**b** Examples of theta frequency and multi-unit activity population averaged firing rate (5 ms average) from electrode sites measured from organoid L2 and L1 respectively. **c** The time that theta oscillations remain above the noise floor increases during population bursts. For each array (at 7 months old), the number of channels that had a correlation score of at least 0.15 with the top correlated reference 'seed' channel was selected. From these channels, the number of channels with at least a 10% increase in the fraction of time that the theta envelope exceeds 1.5 rms for burst periods vs non-burst periods is plotted (blue) and the number of channels with at least a 10% decrease is plotted (red). In all cases, the number of channels with at least a 10% increase is larger, indicating that there are more channels with a higher amplitude during burst periods versus non-burst periods than there are channels with a lower amplitude during burst periods versus non-burst periods for $n = 4$ organoids (organoid L4 and L6 were omitted due to low population rate activity compared to the other organoids).

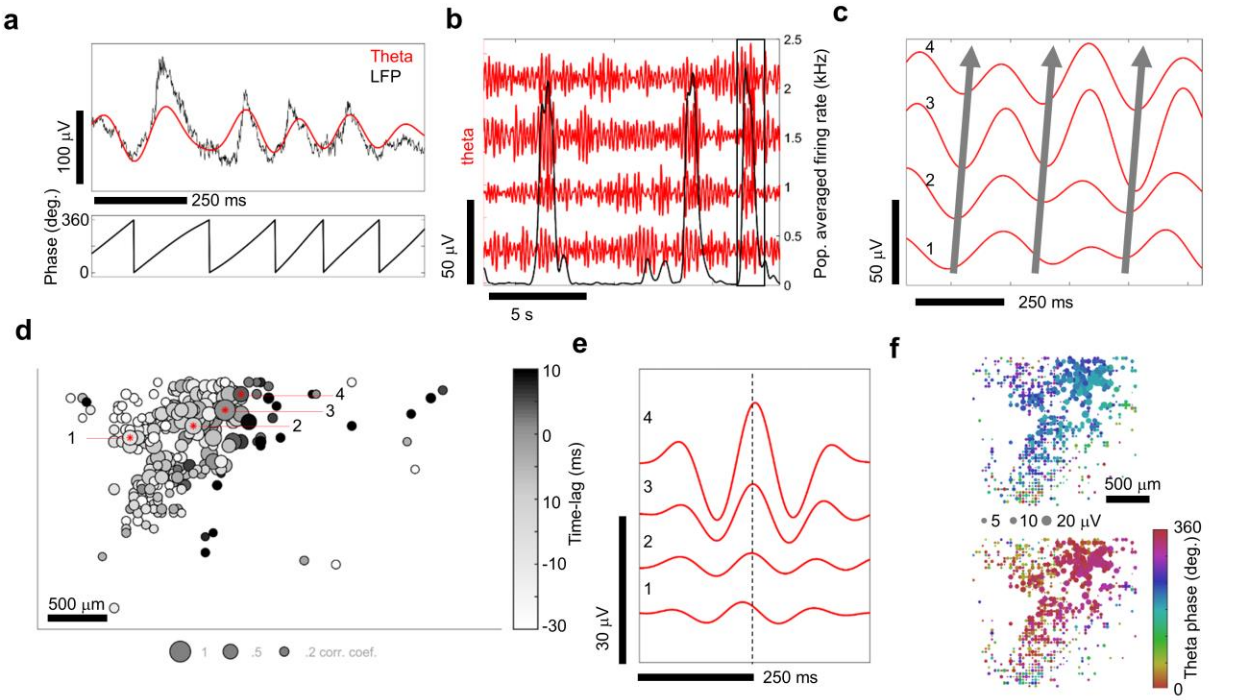


**Supplementary Fig. 17. Coherence of theta oscillations and synchronization with neuronal population bursts.** Additional example of correlated theta activity for organoid L2 at 8 months. **a** The raw local field potential (< 500 Hz, black line) and the 4-8 Hz theta filtered band (red) (top). The phase of the theta oscillation (bottom). **b** Theta band oscillations (red) from four different recording sites and the population-averaged firing rate (black, 100 ms average). **c** Zoomed in view of highlighted gray rectangle in **b**. The solid lines indicate the relative phase offsets of theta oscillations across spatial sites of the organoid. **d** Spatial correlation map of theta oscillations. The correlation coefficient (bubble size) is shown with respect to the seed reference site (3) and the relative time-lag with respect to the reference electrode is shown in grayscale. **e** Signal averaged theta oscillations using peaks from electrode (3) as a reference. The numbers 1-4 in **c**-**e** all refer to the same set of electrodes. **f** Spatial map of signal averaged theta oscillation phase and amplitude relative to reference electrode number 3. Two time points are shown, one at the center of the reference channel $t_0$ (top) and another 60 ms later (bottom).

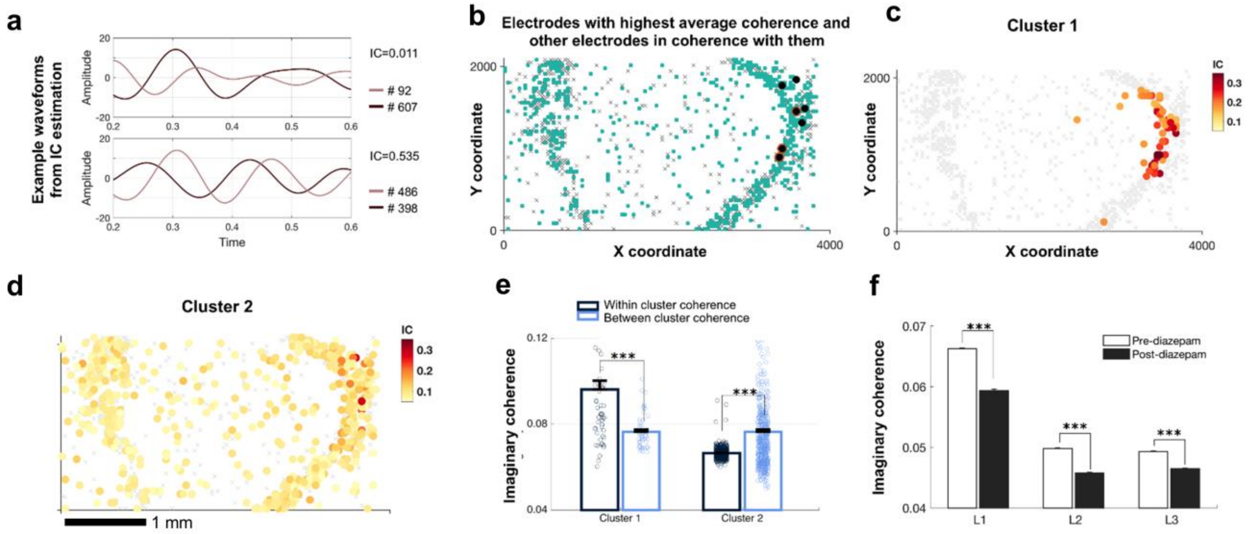

**Supplementary Fig. 18. Imaginary coherence. a** Example theta band waveforms from two electrodes that are weakly correlation (top) and highly correlated (bottom) in the organoid (L1) using the imaginary coherence estimation. **b** Seven nodes (in black) identified as the highest coherent nodes. This cluster was identified by first calculating the node-to-node imaginary coherence and thresholding it at the top 10% of the full coherence matrix with a minimum of 200 connections. The top-seven nodes showed connections to a total of 595 nodes (shown in teal). **c-d** Among these 595 nodes, a K-means clustering procedure identified a distinct local cluster (cluster 1, **c**) as separated from the rest of the nodes which showed a lesser degree of coherence (cluster 2, **d**). **e** The local cluster showed significant higher within coherence, depicting higher local connectivity. The rest of the nodes, labeled as cluster 2, showed relatively poor within cluster coherence. **f** The average global imaginary coherence was decreased after treatment with diazepam (50 µM) for three organoids, L1, L2 and L3 ($t$ = 60.36, 94.83, 63.2257 and $P$ < 1e-4, < 1e-4, < 1e-4, for organoids L1, L2 and L3 respectively).

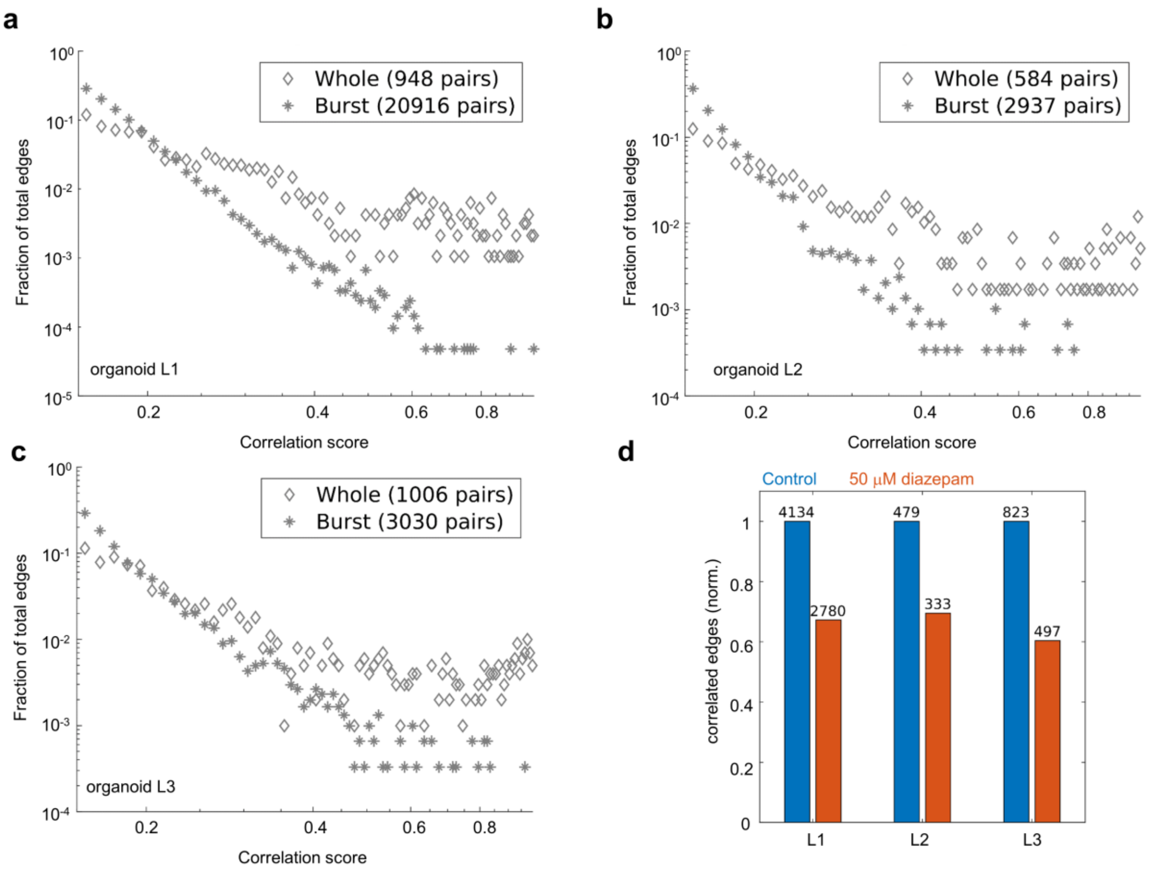


**Supplementary Fig. 19. Distribution of pairwise theta correlations across organoids. a-c** The relative fraction of theta correlation edges (between electrodes) for whole time recording and during burst epochs are shown. The number of correlated edges increases during bursts, but the relative proportion of highly correlated edges that remain outside of bursts increases. **d** The number of pairwise theta correlations between electrode sites (above 0.2) decreases for diazepam (orange) compared to control (blue).

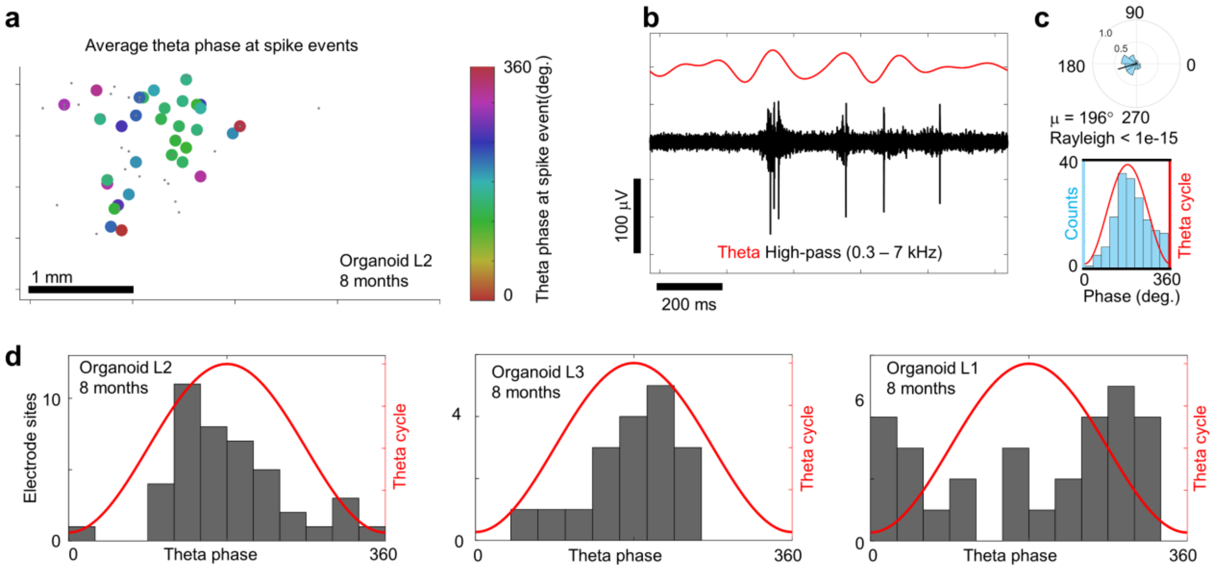


**Supplementary Fig. 20. Distribution of phase-locking of spikes to theta oscillations across multiple organoids. a** Spatial plot of phase-locked spikes to theta oscillation that satisfy the Rayleigh criterion for non-uniformity ($P < 0.05$) for organoid L2 at 8 months. The color indicated the mean theta phase angle (μ) associated with spike events at a given electrode. The gray dots are spiking channels that do not satisfy the Rayleigh criterion ($P > 0.05$). **b** Theta band (4-8 Hz) is shown in red while the spikes are shown in black in the high pass band (0.3-7 kHz) from an individual electrode from **a**. **c** Top, circular distribution of theta-spike angles measured from the electrodes in **b** over three-minute duration. Bottom, distributions of theta-spike angles shown relative to the theta cycle are visualized as a histogram plot. **d** Histogram of the mean phased-locked angle to theta for all electrode sites across the array that satisfy the Rayleigh criteria ($P < 0.05$) for $n = 3$ organoids.
